# Gut-Liver physiomimetics reveal paradoxical modulation of IBD-related inflammation by short-chain fatty acids

**DOI:** 10.1101/706812

**Authors:** Martin Trapecar, Catherine Communal, Jason Velazquez, Christian Alexander Maass, Yu-Ja Huang, Kirsten Schneider, Charles W. Wright, George Eng, Omer Yilmaz, David Trumper, Linda G. Griffith

## Abstract

Association between the microbiome, IBD and liver diseases are known, yet cause and effect remain elusive. By connecting human microphysiological systems of the gut, liver and circulating Treg/Th17 cells, we modeled progression of ulcerative colitis (UC) ex vivo. We show that microbiome-derived short-chain fatty acids (SCFA) may either improve or worsen disease severity, depending on the activation state of CD4 T cells. Employing multiomics, we found SCFA increased production of ketone bodies, glycolysis and lipogenesis, while markedly reducing innate immune activation of the UC gut. However, during acute T cell-mediated inflammation, SCFA exacerbated CD4^+^ T cell effector function, partially through metabolic reprograming, leading to gut barrier disruption and hepatic injury. These paradoxical findings underscore the emerging utility of human physiomimetic technology in combination with systems immunology to study causality and the fundamental entanglement of immunity, metabolism and tissue homeostasis.

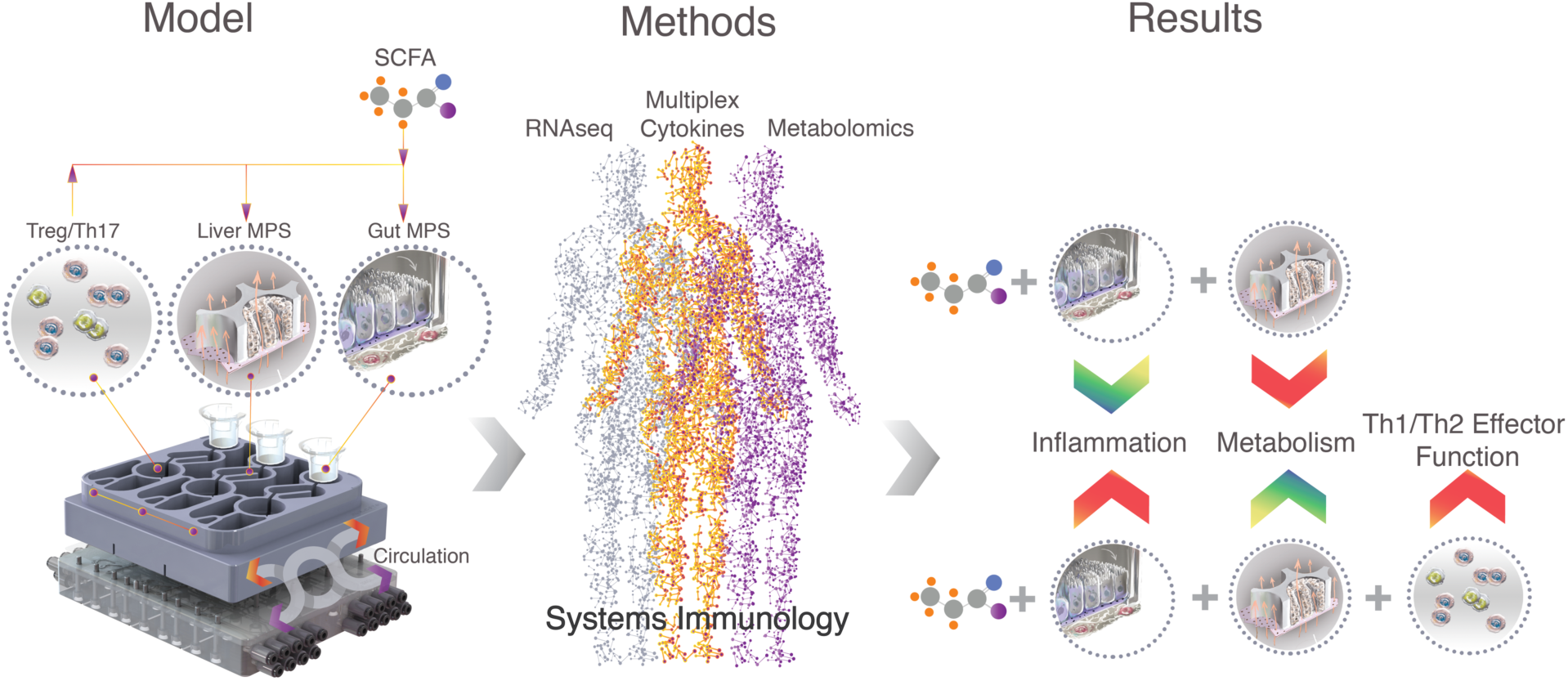

## Introduction

The gut-liver axis is a highly connected system whose roles include processing of gut-derived products, regulation of metabolic homeostasis and stability of systemic immune function (Macdonald and Monteleone, 2005). Thus significant correlations exist between inflammatory bowel disease (IBD), like ulcerative colitis (UC) or Crohn’s disease (CD), and inflammatory liver diseases such as autoimmune hepatitis or primary sclerosing cholangitis (PSC) (Adams and Eksteen, 2006). Risk to develop autoimmune liver disease is increased in patients with IBD and up to 80% of patients with PSC have concurrent UC (Loftus et al., 2005). The manifestations of these interconnected pathologies are relatively well described, yet the “cause and effect” paradox remains to be solved. This is in part due to the overbearing complexity of animal models offering poor control of experimental parameters (Yissachar et al., 2017) and on the other hand lack of *in vitro* models that would take into account the complex organ-organ dynamics needed for accurate representation of organ function (Chen et al., 2017).

A common feature of IBD and extra-intestinal complications is the influx of destructive immune cells (Podolsky, 1991). Most forms of hepatitis show initial infiltration of T cells via the portal vessels and are crucial for the development of chronic liver inflammation (Desmet et al., 1994). At the heart of the balance between autoimmunity and tolerance lie CD4 regulatory T cells (Tregs), producing TGF-β (Rudensky, 2011), and Th17 cells (Th17s) releasing the cytokines IL-17 and IL-22 (Ivanov et al., 2009). Treg/Th17 disbalance has been extensively reported in both IBD (Eastaff-Leung et al., 2010; Zhou and Sonnenberg, 2018), autoimmune liver diseases (Feng et al., 2017) as well as organ rejection (Wu et al., 2017). Furthermore, plasticity of both Tregs and Th17s allows them to transition into more inflammatory CD4 Th1/Th2 hybrid subtypes under acute conditions via antigen and antigen-independent activation. UC is associated with a Th2/ Th17 phenotype and ample production of IL-13, while CD is connected with Th1/Th17 effector response (Fuss et al., 1996; Hoving, 2018; Lavoie et al., 2019).

While many drivers of autoimmunity and inflammation in the gut-liver axis remain unknown, they can be influenced by the microbiome. Short-chain fatty acids (SCFA) are metabolites derived from microbial fermentation of fiber in the colon (Koh et al., 2016). The most abundant SCFA are acetate, propionate and butyrate and their concentration (up to 120 mM total SCFA) is linked to SCFA producing bacterial phyla (Ananthakrishnan et al., 2013; Sampson et al., 2016). SCFA have been shown to exert a variety of beneficial effects on the host (Chang et al., 2014; Koh et al., 2016). Among these are increased gut barrier function, increased liver metabolism, reduction of IBD associated symptoms and promotion of immune tolerance. Contrary to the latter findings, accumulating data shows SCFA to enhance both CD4 (Kespohl et al., 2017; Park et al., 2016; Park et al., 2015) and CD8 (Luu et al., 2018; Trompette et al., 2018) T cell effector function. Several animal and human trials found SCFA to either lack any therapeutic efficacy or to exacerbate inflammation (Ananthakrishnan et al., 2013; Berndt et al., 2012; Breuer et al., 1997; Sampson et al., 2016; Tarrerias et al., 2002; West et al., 2013). Conflicting results suggest the function of SCFA to be context and disease severity dependent, thus their role in T cell activation and metabolism under conditions of gut-liver inflammation in humans demands urgent clarification.

*In vitro* physiomimetic technology, in combination with systems immunology, offers new possibilities for illuminating mechanisms in multi-organ inflammatory diseases by modeling complex interactions with multiple interconnected human microphysiological systems (MPSs) (Ronaldson-Bouchard and Vunjak-Novakovic, 2018; Yissachar et al., 2017). MPSs are *in vitro* models that are both complex - comprising multiple cell types, specialized microenvironments, and perfusion – and yet reductionist, as they capture only the most salient features of *in vivo* organ behavior required to model the physiological process in question (Chen et al., 2017; Edington et al., 2018; Tsamandouras et al., 2017).

Here, we describe a human physiomimetic model of ulcerative colitis that supports integrated co-culture of 3 fluidically-communicating MPSs, featuring individually-addressable on-board microfluidic pumps compatible with highly lipophilic compounds such as steroid hormones.

We used the system to model the gut-liver-immune axis by connecting a gut MPS of UC (primary human UC epithelium, dendritic cells and macrophages) with a liver MPS (healthy human hepatocytes and Kupffer cells) and circulating Treg/Th17 cells. By measuring gene transcription, cytokines and metabolomic changes during different modes of interaction, we gained new insight on the connectedness between UC, liver function and SCFA in an all-human model. While SCFA may reduce innate immune activation of primary tissue, they may also increase the effector function of activated CD4 T cells and thus exacerbate acute UC inflammation and raise the risk of liver injury.

## Results

### Gut MPS of ulcerative colitis donor, directs increased polarization of Treg/Th17 cells into Th1/Th2 cytokine producing effector subtypes in absence of SCFA

First, in the absence of SCFA and under static conditions off-platform, we compared baseline behaviors of healthy and UC gut MPSs on the basis of disease status, immune activation, and growth characteristics through (i) differential gene expression analysis of the epithelia; (ii) immunofluorescent imaging; and (iii) cytokine production and concomitant ability to disrupt Treg/Th17 balance. Gut MPSs were established by seeding equal numbers of epithelial cells from a UC donor and from a non-diseased control on separate Transwell membranes, and then attaching peripheral blood mononuclear cells (PBMC)-derived macrophages and dendritic cells on the basolateral side of the membrane after monolayer differentiation. All experiments were conducted in a slight modification of culture medium, except for the apical gut medium, which had previously been tailored for physiological inflammatory responses of the human liver MPS, in which the extreme supra-physiological concentrations of cortisol and insulin typically used to maintain CYP450 levels in primary hepatocyte cultures were reduced to concentrations within the physiological range (Tsamandouras et al., 2017).

After 4 days of epithelial co-culture with innate immune cells, gene expression in epithelia harvested from the apical side of replicate Transwell co-cultures showed classic hallmarks of UC disease phenotype when UC epithelia were compared to the control epithelia using pathway enrichment based on GEO (Fig.1A). Further, gene ontology (GO) enrichment analysis (Fig.1B) showed a strong upregulation of pathways associated with innate immune activation, apoptotic signaling and EGF activity in the UC epithelia. Pathway analysis based on KEGG (Fig.1C) revealed increased cancer-related signaling through MAPK and PI3K-Akt pathways, which indicates increased cancerous neoplasticity and possible progression into carcinoma. Morphologically, epithelia from both donors formed confluent monolayers (Fig.1D) and exhibited comparable transepithelial resistance (Fig.1E). However, monolayers of UC, but not healthy gut MPSs, showed enlargement of hyperchromatic nuclei and irregular actin fragmentation, indicators of IBD dysplasia (Fig.1E) (DeRoche et al., 2014). These morphological findings are consistent with the observation that differentially-expressed genes (Fig.1B,C) include those associated with regulation of actin, focal adhesions and cell junction, which is indicative of IBD-associated remodeling (Kinchen et al., 2018).

**Fig.1.**
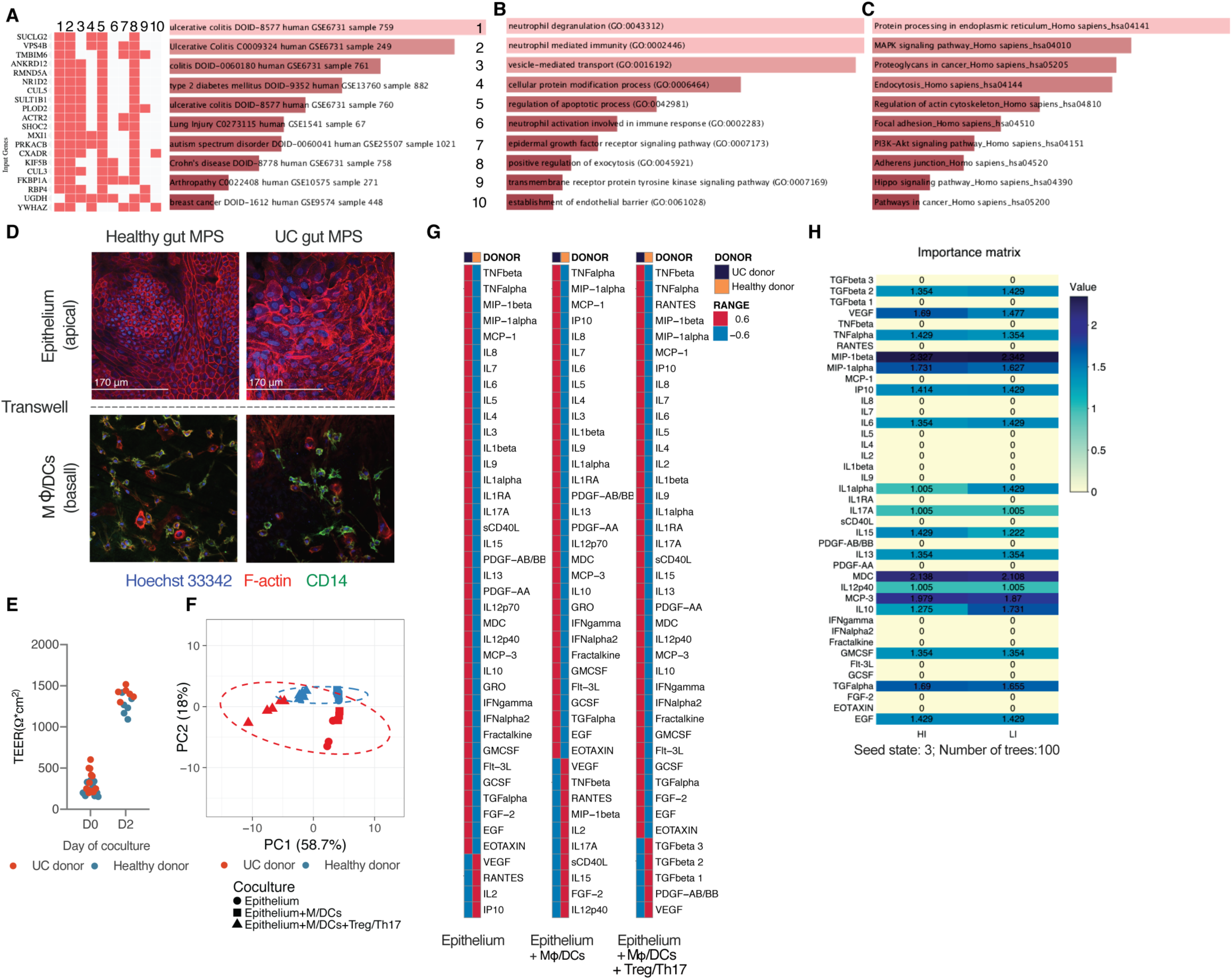
Co-culture of the UC gut MPS with Treg/Th17 cells leads to increased production of Th1 and Th2 inflammatory cytokines. Differential gene expression pathway enrichments in UC gut MPS as compared to healthy control MPS based on GEO disease related pathway enrichments (A), GO Biological process terms (B) or KEGG (C). (A-C) Data represent averages of 3 biological replicates. (D) Representative confocal images of gut MPS from the healthy gut MPS (left) and UC gut MPS (right). Top: epithelial monolayers seeded on the apical side of transwell membranes. Bottom: Co-cultured macrophages and dendritic cells seeded on the basal side of transwell membranes (E) Representative TEER values of UC and healthy gut MPS at day 0 of gut MPS development and at day 2. (F) Representative PCA of multiplexed cytokines/chemokines released by each donor’s epithelium alone, their corresponding MPS, and MPS interacting with Treg/Th17 cells. (G) Collapsed heatmaps based on means representative of 3-5 biological replicates indicating Z scores of cytokines/chemokines expressed under each condition as described under (F). (H) Importance matrix derived from random forest analysis of cytokines/chemokines released by gut MPSs in co-culture with Treg/Th17s. For actual concentration values of individual cytokines/chemokines measured consult Figure S1.

We next compared the inflammatory potential of the UC vs. control donors on the basis of cytokine production by the individual epithelium alone, epithelium in co-culture with macrophages/dendritic cells (i.e. gut MPS) and during gut MPS co-culture with PBMC derived Tregs and Th17 cells (ratio of 2:1) from the same donor as the antigen-presenting cells.

Analysis of 41 cytokine concentrations in the basal medium after 48 h by principal component analysis clearly distinguishes individual donors in all 3 conditions based on cytokine/chemokine production (Fig.1F), where 59% of the variability was explained by the presence or absence of Treg/Th17 cells and 18% by the donor. Mean concentration values of the majority of measured cytokines/chemokines were more highly expressed by the UC donor regardless of the co-culture condition (Fig.1G, Fig. S1A,B). Interestingly, VEGF concentrations were persistently higher in all conditions with the healthy donor despite VEGF’s known association with UC. This can be explained by a higher expression of VEGF receptors on UC epithelium and increased receptor-mediated VEGF internalization and degradation (Fig.S1C) thus reducing measurable free autocrine VEGF (Barr et al., 2012). Despite twice the number of Tregs over the number of Th17s, CD4^+^ T cells began to polarize in their cytokine production toward a mix of Th1, Th2, Th17 cytokine producing cells in co-culture with the UC gut MPS. The UC donor MPS induced T cell production of IL-4, Il-5, IL-13, TNF-α and interferons (Fig.S1A). This set of increased inflammatory cytokines coincides with previous reports showing UC epithelia to predominantly induce Th2/Th17 conditions (Fuss et al., 1996). The Treg/Th17 balance remained intact in co-culture with the healthy gut MPSs as indicated by the preserved Treg production of TGF-β1 and -β2 and low IL-17A release (Fig. S1B). A study using in vitro models of the intestinal organ system, albeit that of mice and in absence of other organ systems, showed epithelial cultures exposed to different microbes to dictate Treg/Th17 polarization (Yissachar et al., 2017). The machine learning algorithm Random Forests was able to predict the MPS donor with 100% accuracy when co-cultured with Treg/Th17 cells (Fig.S1D); the most predictive factors were the pleiotropic lymphocyte attractants MIP-1β, MDC, and MCP-3 (Fig.1F). Both epithelial donors share the HLA alleles HLA A*02, B*51, DQA1*04, DQB1*06 and DRB1*03 which taken together with the physical separation of the epithelium and co-cultured underlying immune cells via 0.4µm microporous membranes, makes the rapid activation of CD4 T cells due to greater donor-mismatch highly unlikely.

Throughout the rest of the study, the UC gut MPS was used to further clarify the interaction of SCFA, inflamed gut tissue, liver MPS, and circulating T cells.

### Bioaccessible SCFA lead to enrichment in metabolic pathways and reduction of inflammatory pathways in the UC gut MPS

As influence of SCFA on healthy epithelium has been described before (Koh et al., 2016), we focused on investigating its effects on the inflamed UC tissue and MPS. First, we investigated the interaction of SCFA and UC epithelium under static conditions alone without the presence of antigen presenting cells. Acetate, propionate and butyrate in a physiological molar ratio of 6:2:2 (Cummings et al., 1987) at a total combined concentration of 20mM were added to the apical media of UC epithelial monolayers in transwell inserts. After 4 days of culture (with a media and SCFA change at 48 hr), tissue was harvested, RNAseq performed, and differential gene expressions were established against untreated control groups (Fig.2 A,B). KEGG pathway enrichment analysis showed a significant upregulation of Foxo and the PPAR signaling pathway (Fig. 2B), mainly PPAR-α/δ, which drives glycerolipid metabolism upon fatty acid stimulation and ketone body production (Puchalska and Crawford, 2017). Concurrently, cell cycle pathways were down regulated as well as genes associated with proteoglycan synthesis in cancer. In vitro studies have shown SCFA to inhibit proliferation of cancer cell lines in addition reduction in butyrate-producing bacteria is linked to both colorectal cancer development and progression (O’Keefe, 2016).

**Fig.2.**
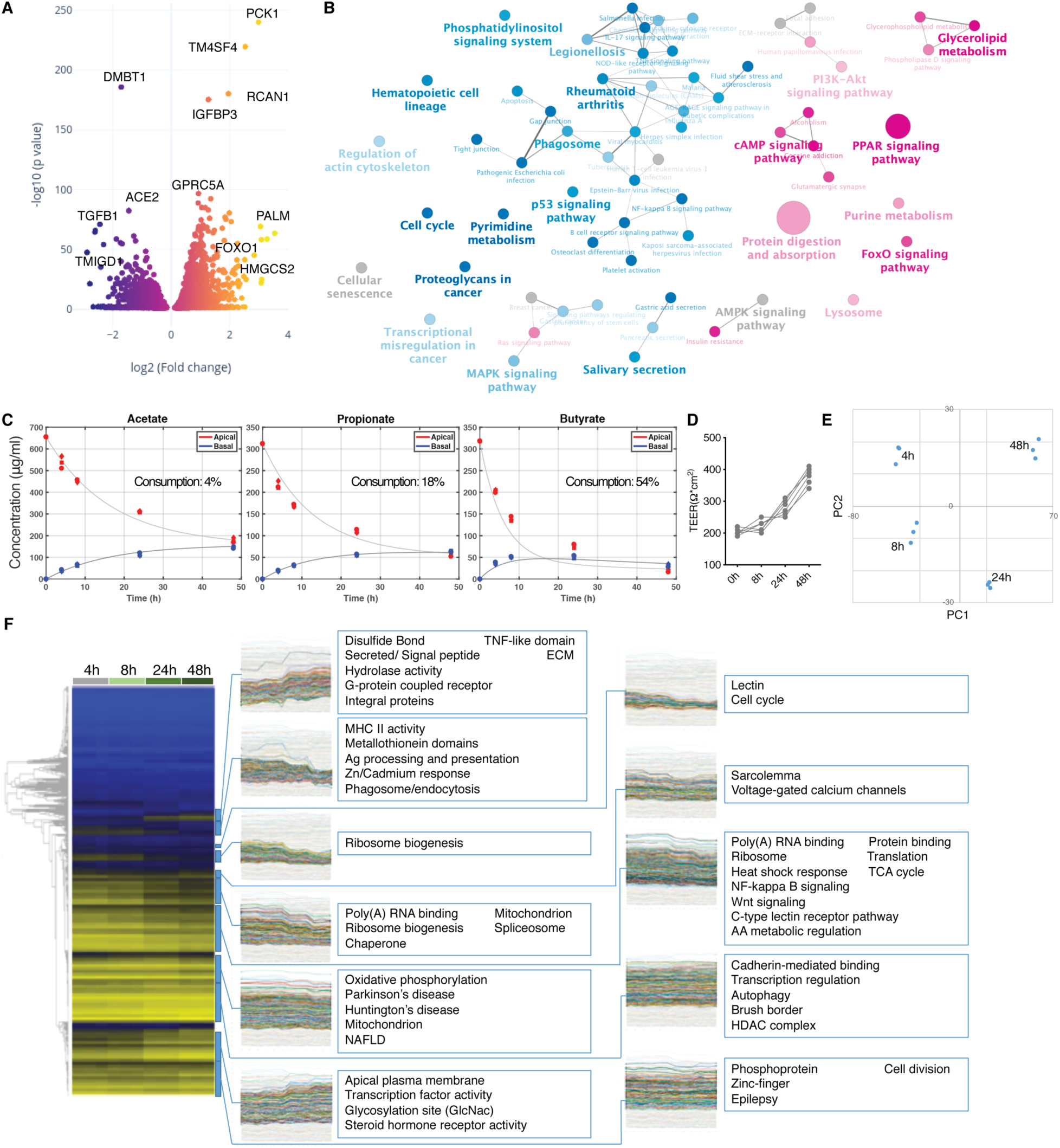
Bioaccessible SCFA affect gene expression of UC epithelium and the UC gut MPSs. (A) Volcano plot of differentially expressed genes (DEG) in monolayers of UC epithelium with SCFA over controls without. (B) Network of enriched (magenta) and suppressed (blue) KEGG pathways based on DEG shown under (A). (C) Apical and basal concentrations of acetate, propionate and butyrate over 48h in UC gut MPSs upon an initial apical total SCFA concentration of 20mM. Consumption of each SCFA by the epithelium is shown as percent loss in corresponding plots. Each time point was measured in 3 biological replicates. (D) Representative TEER values of gut MPSs cultured with 20mM of SCFA. (E) PCA analysis of changes in epithelial gene expressions at each time point shown under (C). (F) Epithelial tissue from each time point shown under (C) was harvested for RNAseq analysis. Left: Heat-maps with gene-level dendrograms. Right: Clades from the gene-level hierarchical clustering with gene expression time-course representations of each gut MPS interacting with SCFA. Each time point represents 3 biological replicates.

Next, we established the bioaccessibility and clearance of SCFA (20mM) by the UC gut MPS with macrophages and dendritic cells under basolateral circulation at a flow rate of 1µL/s. Over the course of 48 hours, 54% of butyrate was consumed by the UC gut MPS while the majority of acetate (96%) and propionate (82%) passed the gut barrier into the basal compartment (Fig.2C). These results correlate with human clinical data that show butyrate serves as the primary energy source of colonocytes and that in average only 50% of butyrate reaches the portal vein (Cummings et al., 1987). As a result, the molar ratio of SCFA in portal blood switches to 7:2:1, which mirrors our findings. Epithelial barrier was maintained throughout the time course (Fig.2D). Concurrently, epithelial tissue was harvested at 4 h, 8 h, 24 h and 48 h post SCFA application to identify time dependent changes in gene expression and to validate SCFA effects seen under Figure 2B. Principal component analysis of expressed genes showed significant changes throughout the time course (Fig.2E). Further analysis repeatedly indicated downregulation of UC gut MPS genes governing cell cycle and WNT signaling through FoxO as reported *in vivo* (Kaiko et al., 2016), as well as upregulation of hormone receptor activity, expression of integral membrane proteins and ECM remodeling. Moreover, in the coculture of the inflamed UC epithelium with antigen-presenting cells, SCFA downregulated inflammatory NF-kappa B signaling pathways as seen with the epithelium alone (Fig.2B), which confirms previous reports on anti-inflammatory potential of SCFA (Koh et al., 2016).

### Interaction between SCFA, gut and liver MPSs results in decreased innate inflammation of the gut MPS and increased hepatic metabolism

Having established that apical SCFA are transported across the colonic epithelial barrier, partially metabolized by the colonic epithelia, and reduce signatures of inflammation of UC epithelia compared to controls without SCFA, we next used our physiomimetic platform, the 3XGL (3 modules of “gut-liver” per platform; Fig.S2A), to investigate the effects of SCFA on the gut-liver axis. We focused initial experiments on behaviors of the gut-liver axis in the absence and presence of SCFA without adaptive immune cells, using the gut and liver MPSs connected at recirculating flow rates of 30 ml/day. Integration of gut and liver MPSs significantly affects their individual phenotypes and gene expression (Fig.3A,B) in accordance with our previous observations during interaction of a CaCo-2 cell based MPS with the liver MPS (Chen et al., 2017; Tsamandouras et al., 2017). The most notable effects of interaction on the gut MPS in our study were increased pentose and glucuronate interconversions, ABC transporters and protein digestion as well as a decrease in colorectal cancer related pathways (Fig.3A). In the same time interaction between the gut and liver MPSs lead to increased CYP450 activity, biosynthesis of unsaturated fatty acids and steroid hormones with downregulation of inflammatory pathways in the liver MPSs (Fig.3B).

**Fig.3.**
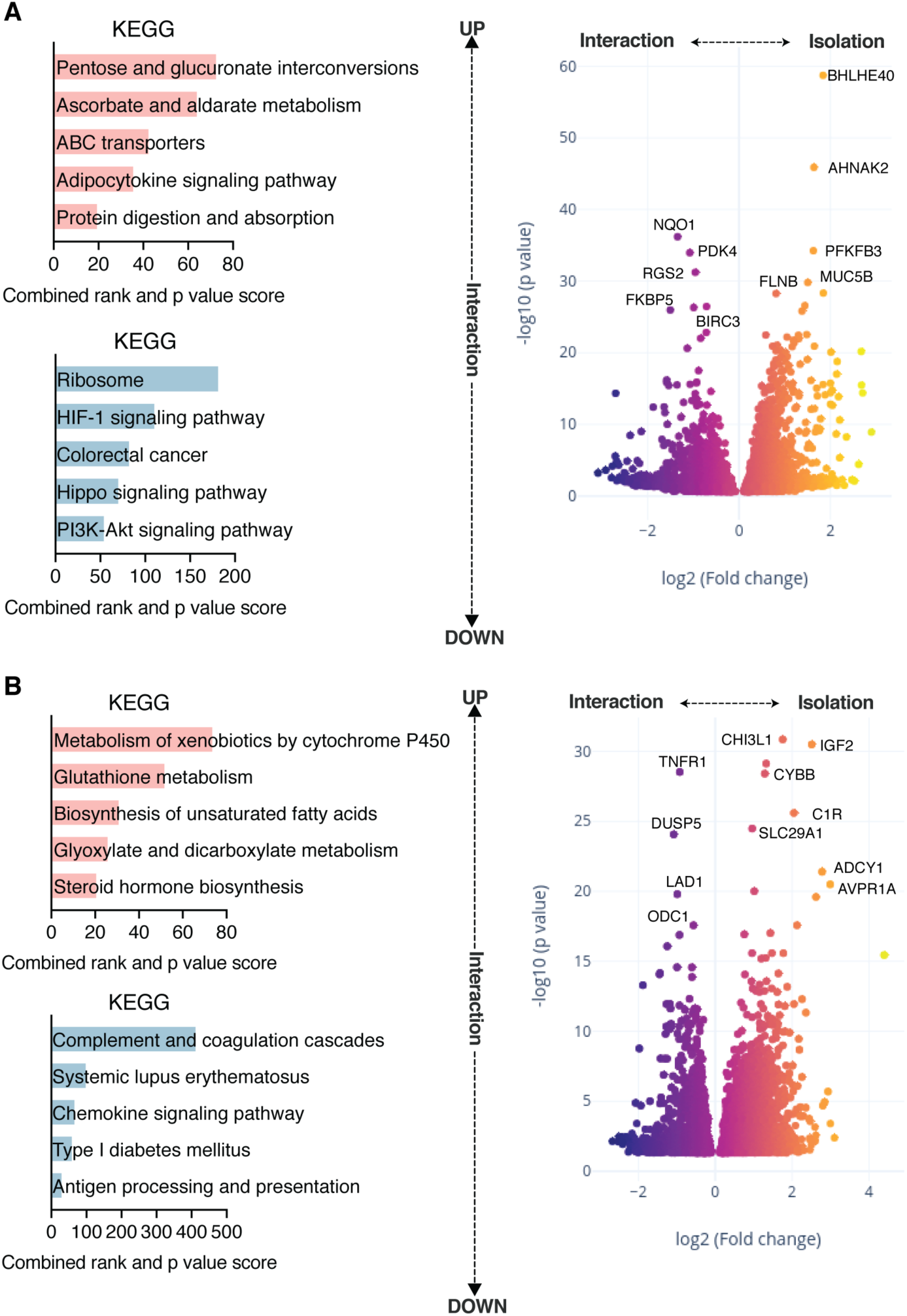
The integration of UC gut and liver MPS significantly affects their individual phenotype and gene expression. (A) DEG in UC gut MPS in isolation or in interaction with the liver MPS was assessed. Left: Upregulated (top) and downregulated (bottom) pathway enrichments based on KEGG in the gut MPS during interaction with the liver MPS. Right: Volcano plot showing DEG comparing gut MPSs in interaction with the liver MPS or in isolation. (B) DEG in liver MPS in isolation or in interaction with the UC gut MPS. Left: Upregulated (top) and downregulated (bottom) pathway enrichments based on KEGG in the liver MPS during interaction with MPS of the gut. Right: Volcano plot showing DEG comparing liver MPSs in interaction with the gut MPS or in isolation. Data represents 3 biological replicates.

We next performed a 4-day interaction experiment between gut and liver MPSs with the addition of 20 mM of total SCFA into the apical gut media. On day 4 (after an interim medium and SCFA refreshment on day 2), media were collected for cytokine and metabolomic analyses and tissue harvested for RNAseq (Fig.4A-H). During interaction, SCFA were both adsorbed and metabolized by the gut MPS and further metabolized by the liver MPS (Fig.S3A) at similar rates when measured in day 2 as well as day 4 media. Liver clearance rates for SCFA, derived with PBPK models using concentrations of SCFA measured in the gut basal compartment during interaction (Fig. S3A) and gut MPS absorption and metabolism data in isolation (Fig.2C), show a pattern consistent with known in vivo clearance: near-complete clearance of butyrate, intermediate clearance of propionate, and considerably lower clearance of acetate. Gene expression clustering clearly separates the gut MPS epithelial tissue from liver MPS samples and validates the presence of tissue specific genes (Fig.S3B). SCFA induced significant changes in both MPS compared to control interactions (Fig.4A,C). During gut-liver interaction, changes in UC tissue upon addition of SCFA are highly consistent with changes observed in isolation studies: upregulation of PPAR, hormone synthesis pathways and suppression of NF-kappa B signaling based on KEGG was observed (Fig.4B). Moreover, pathways of the glutaminergic synapse were enriched after addition of SCFA due to increased expressions of glutamate metabolism genes with implied neurotropic effects of SCFA. Epithelial TEER measurements of both gut-liver and gut-liver-SCFA interaction groups were comparable to the control in isolation and were maintained in the in vivo physiological range between 100-400 Ohms/cm^2^ (Srinivasan et al., 2015). Hepatic albumin production increased throughout this interaction (Fig.S3C,D,E). SCFA added to the apical compartment of the gut MPS, absorbed, and transported to liver MPS induced upregulation of several metabolic pathways of the liver MPS in accordance with observations in vivo (Morrison and Preston, 2016), including gluconeogenesis, retinol and tyrosine metabolism as well as bile acid synthesis and secretion, as confirmed by KEGG analysis (Fig.4D). Decreased innate activation of the UC gut MPS had an anti-inflammatory effect on the liver MPS, linking the state of gut activation to liver homeostasis. UC is tightly connected to liver PSC, and clinical data show that even after liver transplantation, UC and gut inflammation pose an increased risk for PSC recurrence (Cholongitas et al., 2008).

**Fig.4.**
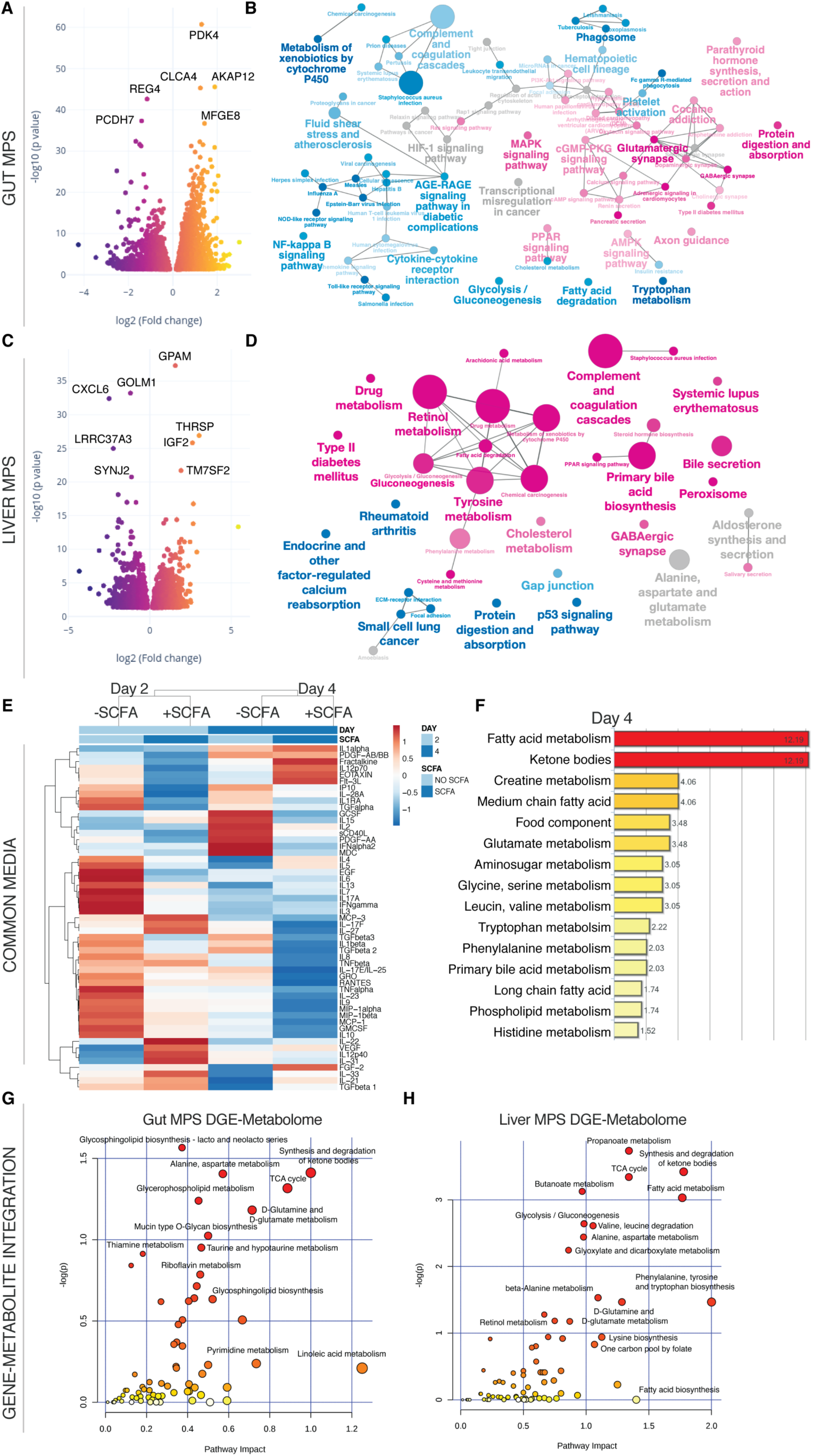
SCFA reduce innate immunity of the UC gut MPS and beneficially affect hepatic metabolism during interaction. (A,C) Volcano plots of DEG in the gut MPS (A) and the liver MPS (C) in presence or absence of SCFA. (B,D) Network of enriched (magenta) and downregulated (blue) pathways based on KEGG in gut MPS (B) or liver MPS (D). (E) Clustered heatmap of multiplexed cytokines/chemokines present in circulating common media at day 2 and 4 during gut-liver MPS interaction +/- SCFA. For concrete concentrations of measured proteins and their importance consult Fig.S4. (F) Significantly enriched metabolic pathways based on quantification of metabolites found in the common medium at day 4 of gut-liver-SCFA interactions. (G,H) Integration of significantly upregulated genes in the UC gut MPS (G) or liver MPS (H) during interaction with SCFA and enriched metabolites found in the common medium at day 4. Pathway impact score is shown on the x-axis and statistical significance on the y-axis. All data represents 3 biological replicates.

Addition of SCFA to the apical gut MPSs in the gut-liver interaction condition had a dramatic effect on production of cytokines released into the common medium. Cytokine analysis of the shared common medium showed significant reduction of inflammatory mediators in the SCFA treated group vs the non-treated group (Fig.4E, Fig.S4A,B). Most predictive reductions distinguishing these two groups were in PDGF, IL-9, GRO and MDC expression based on random forest analysis (Fig.S4C). Reduced secretion of the chemoattractants GRO and MDC correlates to decreased enrichment of innate immune pathways.

Global metabolomic discovery of the common media from samples collected at day 4 for the +/- SCFA conditions in gut-liver interaction indicated enrichment in fatty acid metabolism, ketone body synthesis and increased creatine metabolism (Fig.4F) related to increased PPAR signaling in the presence of SCFA (Puchalska and Crawford, 2017), consistent with gene expression results for UC epithelia in isolation (Fig.2B). Glucose consumption and conversion into lactate was higher in the gut-liver-SCFA interaction at day 2 but not at day 4 (Fig.S5A) which implies early increased global metabolic activity supported by the increase of TCA intermediate aconitate (Fig.S5B). Ketone bodies are produced by the liver under fasting condition through conversion of SCFA and by conversion of butyrate in the colon (Henning and Hird, 1972; Kimura et al., 2011). In our system, production of ketone bodies without SCFA stems predominantly from the liver MPS and is increased by SCFA (Fig.S5C). The gene-metabolite integration of overlapping gene and metabolomic enrichments for each MPS reveals common as well as individual metabolic pathways of both tissues (Fig.4G,H). SCFA induced glycerophospholipid and glycersphyngolipid metabolism are more prevalent in the gut MPS. They are tightly connected to gut health by promotion of cytokine synthesis and cell structure integrity (Yan et al., 2016). Fatty acid, butanoate and propionate metabolism was highly induced in the liver MPS confirming liver metabolism of these SCFA.

### Inclusion of Treg/Th17 cells during gut-liver interaction leads to UC triggered acute T cell mediated inflammation and autoimmune hepatitis

We introduced Treg/Th17 cells in a 2:1 ratio and equal numbers into all compartments of the platform with circulation-enabled transport between MPSs. Gene cluster analysis indicated partial engraftment of Treg/Th17 cells within the liver tissue itself based on the expression of FOXP3, ICOS and CD28 (Fig.S3B). As observed in studies of gut MPSs in isolation, the UC gut MPSs induced a high degree of activation of added CD4 T cells, which led to decreased gut barrier function and reduced liver albumin secretion (Fig.S4C,D,E) . In parallel 4-day interaction studies where the same donors of liver hepatocytes, Kupffer cells and PBMCs were used as in this experiment, but with the characterized healthy epithelium donor, Th1/Th2 activation was largely absent and gut barrier as well as liver albumin production preserved (manuscript in preparation).

The CD4 T cell-driven inflammation led to significant upregulation of interferon-induced genes in both tissues as compared to the control gut-liver interaction without lymphocytes (Fig.5A,C). Pathways associated with Th1/Th2 differentiation and IBD were highly enriched based on KEGG (Fig.5B) in the gut MPS with increased NOD signaling as well as reduction of lipid metabolism. NOD2 was the first gene associated with IBD and its expression promotes Th17 differentiation (Khor et al., 2011). In response hepatic tissue showed strong enrichment of both Th17 and Th1/Th2 differentiation pathways (Fig.5D). In mice, blocking migration of intestinal effector T cells prevented hepatic allograft rejection establishing a clear link between the gut and autoimmune disease (Murai et al., 2003). A high correlation exists also between UC and autoimmune hepatitis with or without fibrotic PSC (Perdigoto et al., 1992). In mice, T cells were shown to accumulate in the portal region of the liver and drive Th1-like autoimmune liver inflammation (Griffin et al., 2000) while Th17 and Th2 CD4 T cells were associated with tissue fibrosis and injury (Gieseck et al., 2018).

**Fig.5.**
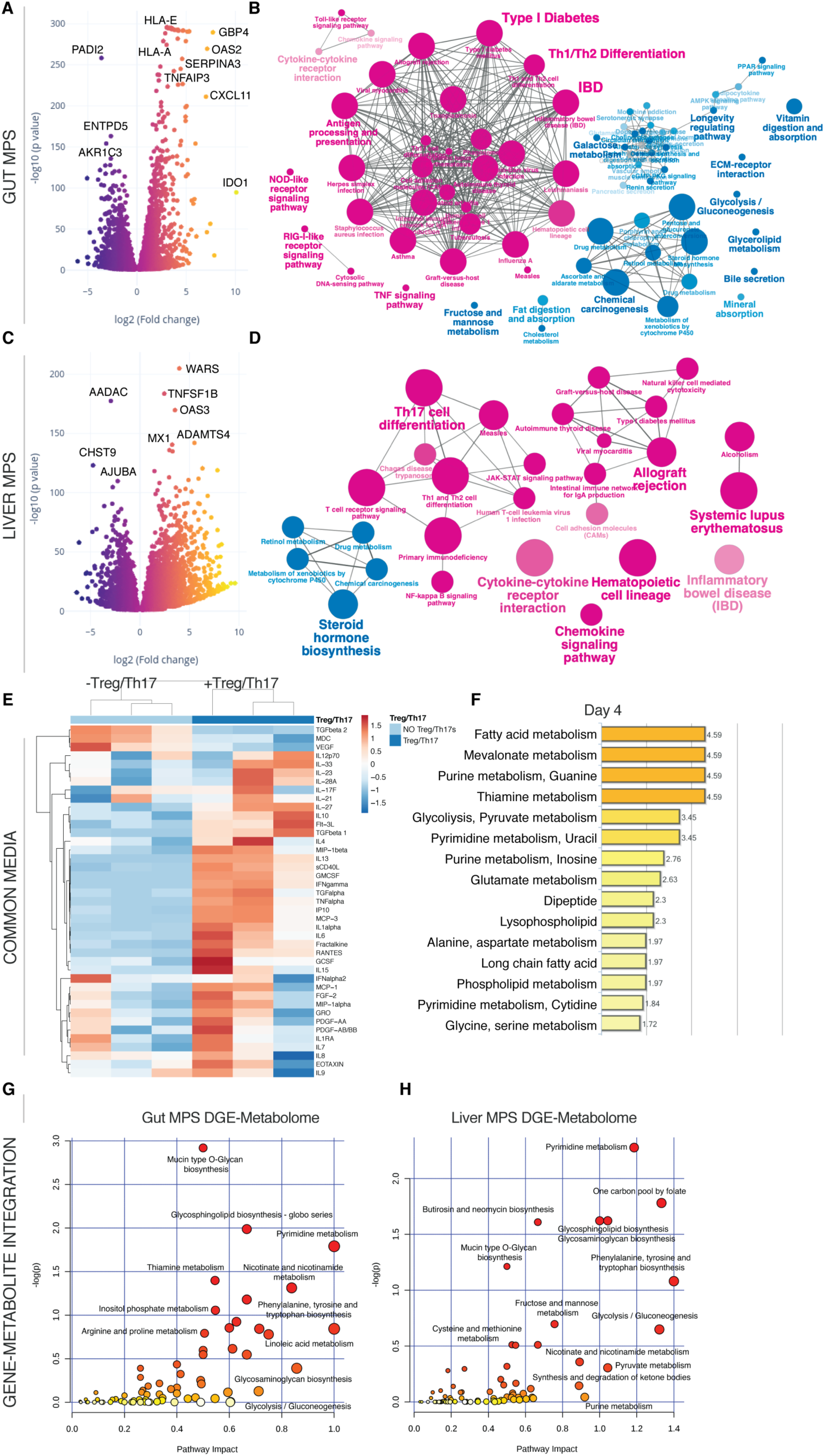
Inclusion of Treg/Th17 cells in the UC gut-liver MPS interaction leads to their conversion into inflammatory Th1/Th2 cells. (A, C) Volcano plots of DEG in the gut MPSs (A) during interaction with the liver MPSs (C) in presence or absence of Treg/Th17 cells. (B, D) Network of enriched (magenta) and downregulated (blue) pathways based on KEGG in gut MPSs (B) or liver MPSs (D). (E) Clustered heatmap of multiplexed cytokines/chemokines present in circulating common media at day 4 during gut-liver MPS interaction +/- Treg/Th17 cells. (F) Significantly enriched metabolic pathways based on quantification of metabolites found in the common medium at day 4 of gut-liver-Treg/Th17 interactions. (G,H) Integration of significantly upregulated genes in the UC gut MPSs (G) or liver MPSs (H) during interaction with Treg/Th17 cells and enriched metabolites found in the common medium at day 4. Pathway impact score is shown on the x-axis and statistical significance on the y-axis. All data are derived from 3 biological replicates.

Predictably, cytokine/chemokine concentration of inflammatory mediators, such as IFN-γ, TNF-α, IL-1α and RANTES, were strongly increased in the presence of activated lymphocytes at day 4 (Fig.5E). While fatty acid metabolism remained intact based on global metabolomic enrichments (Fig.5F), both glycolysis and purine metabolism increased at day 4. Acute inflammation of tissue is associated with a drastic shift in metabolism towards aerobic energy transformation including increased purinergic signaling (Eltzschig et al., 2013) and glucose consumption (Fig.S5A) by T cells. This is in line with the observed enrichment of the mevalonate metabolism at day 4, which is a result of active T cell conversion of glucose to acetyl-CoA due to the Warburg effect (Okin and Medzhitov, 2016). Gene-metabolite integration analysis also indicates strong upregulation of the pyrimidine and nicotinamide metabolism in both tissues (Fig.5G-H). A significant enrichment of sphingolipids was detected, which correlates to clinical data identifying sphingolipids as clinical markers of autoimmune hepatitis in patients’ serum (Lian et al., 2015).

### T cell mediated acute inflammation is exacerbated by SCFA, leading to hepatic failure and gut barrier disruption

As innate immune activation was reduced by SCFA in the inflamed gut MPSs as well as during interaction with the liver MPSs, we next added 20 mM of total SCFA into the apical gut medium in the model of acute CD4 T cell-mediated gut-liver inflammation. Compared to controls without SCFA, inflammation and tissue damage were surprisingly further amplified with their addition. The differential gene expression of the gut MPSs showed asymmetrical downregulation of homeostatic genes and further potentiation of genes related to immune activation (Fig.6A). NF-kappa B with TNF signaling pathways and pathways associated with systemic inflammation were strongly enriched in combination of mTOR activity (Fig.6B) and coinciding with further damage of the epithelial barrier (Fig.S3C,D), indicating irreparable tissue damage. While T cell-mediated inflammation prompted considerable reduction of TEER, addition of SCFA lead to complete barrier failure. Similarly, albumin production of the liver MPS stalled in both groups (Fig.S3E). This observation correlates with wide-ranging downregulation of major metabolic pathways (Fig.6D) and enrichment in MAPK related pathways.

**Fig.6.**
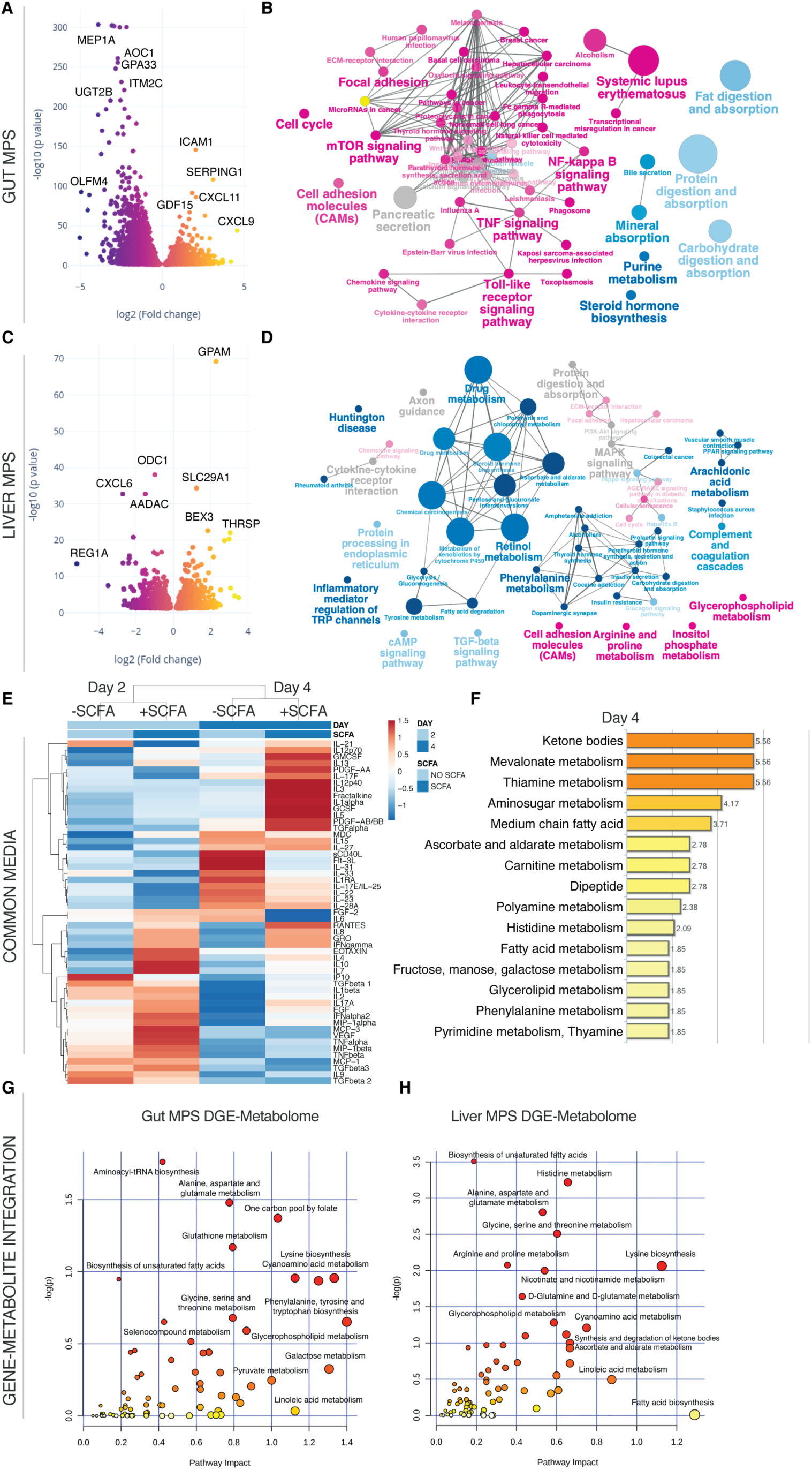
T cell mediated inflammation and tissue injury are significantly exacerbated by SCFA. (A,C) Volcano plots of DEG in the gut MPSs (A) during interaction with the liver MPSs (C) and Treg/Th17 cells in presence or absence of SCFA. (B,D) Network of enriched (magenta) and downregulated (blue) pathways based on KEGG in gut MPSs (B) or liver MPSs (D). (E) Clustered heatmap of multiplexed cytokines/chemokines present in circulating common media at day 2 and 4 during gut-liver MPS interaction +/- SCFA. For actual concentrations of measured proteins and their importance consult Fig.S6. (F) Significantly enriched metabolic pathways based on quantification of metabolites found in the common medium at days 2 and 4 of gut-liver-SCFA interactions. (G,H) Integration of significantly upregulated genes in the UC gut MPSs (G) or liver MPSs (H) during interaction with Treg/Th17 cells and SCFA, and enriched metabolites found in the common medium at day 4. Pathway impact score is shown on the x-axis and statistical significance on the y-axis. All data represents 3 biological replicates.

Multiplex cytokine/chemokine analysis showed increased production of inflammatory cytokines when SCFA were added (Fig.6E, Fig.S6A,B). In particular IFN-γ, IL-5, IL-8, IL-10, and IL-13 were the most condition-predictive cytokines based on random forest analysis (Fig.S6C-E) which is indicative of both Th1 and Th2 expansion. Interestingly, several cytokine concentrations (e.g. IFN-γ and TNF-α) measured during this interaction were similar to reported levels in serum of patients with IBD (Kleiner et al., 2015; Maeda et al., 1992).

In the group with added SCFA, tryptophan conversion into kynurenate as part of the inflammatory IDO pathway was significantly higher as well (Fig.S7A). A similar trend was observed with the kynurenate downstream product quinolinate that serves as precursor of nicotinamide (Fig.S7B). The global metabolomic profile shows enrichments of ketone bodies similar to the gut-liver-SCFA interaction without Treg/Th17 cells (Fig.6F). Mevalonate and thiamine metabolism were further enriched during T cell-mediated inflammation and addition of SCFA. Both glucose consumption and production of pyruvate, lactate and aconitate were highest in the gut-liver-Treg/Th17-SCFA interaction group at day 2 but not day 4, which might be explained by increased tissue failure by that point (Fig.S5A). Increased lactate production indicated a metabolic switch into inflammation-driven aerobic glycolysis. The gut MPS showed enriched glutathione metabolism over the controls without SCFA based on gene-metabolite integration while alanine, aspartate and glutamate metabolism were enriched in both gut and liver MPSs (Fig.6G,H). Glutamate metabolism is essential for T cell function under inflammation and its downstream product glutathione for reduction of oxidative stress (Roth et al., 2002). Furthermore, the liver MPSs showed significant enrichment in metabolism of histidine, a histamine precursor and vital inflammatory agent.

### SCFA act as inhibitors of HDAC and TRAF6 in activated CD4 T cells and stimulate Th1/Th2 differentiation with significant IL-13 production

SCFA have been previously shown in several studies to promote regulatory T cell development in the gut under steady-state conditions (Smith et al., 2013). However, during the gut-liver-SCFA interaction study, we showed SCFA to exacerbate acute inflammation upon the presence of CD4 effector T cells. These contradictory results led us to test the hypothesis that SCFA might affect CD4 T cells differently depending on their state of activation. We therefore performed an isolation study with the same 2:1 ratio of Treg to Th17 cells under steady-state and inflammatory conditions with SCFA. SCFA were added at an average concentration found in the common media (1 mM total) during the interaction study and at the physiological 7:2:1 ratio of acetate, propionate and butyrate. We stimulated human Treg/Th17 cells with IL-12/IFN-γ, which led to upregulation of inflammatory JAK-STAT and TNF pathways as well as to CD4 T cell specific differentiation into Th1/Th2 subtypes (Fig.7A). Next, we determined common and unique gene expressions comparing the 4 different groups; in the group of activated Treg/Th17 cells with SCFA, we identified 308 uniquely downregulated and 724 uniquely upregulated genes (Fig.7B).

**Fig.7.**
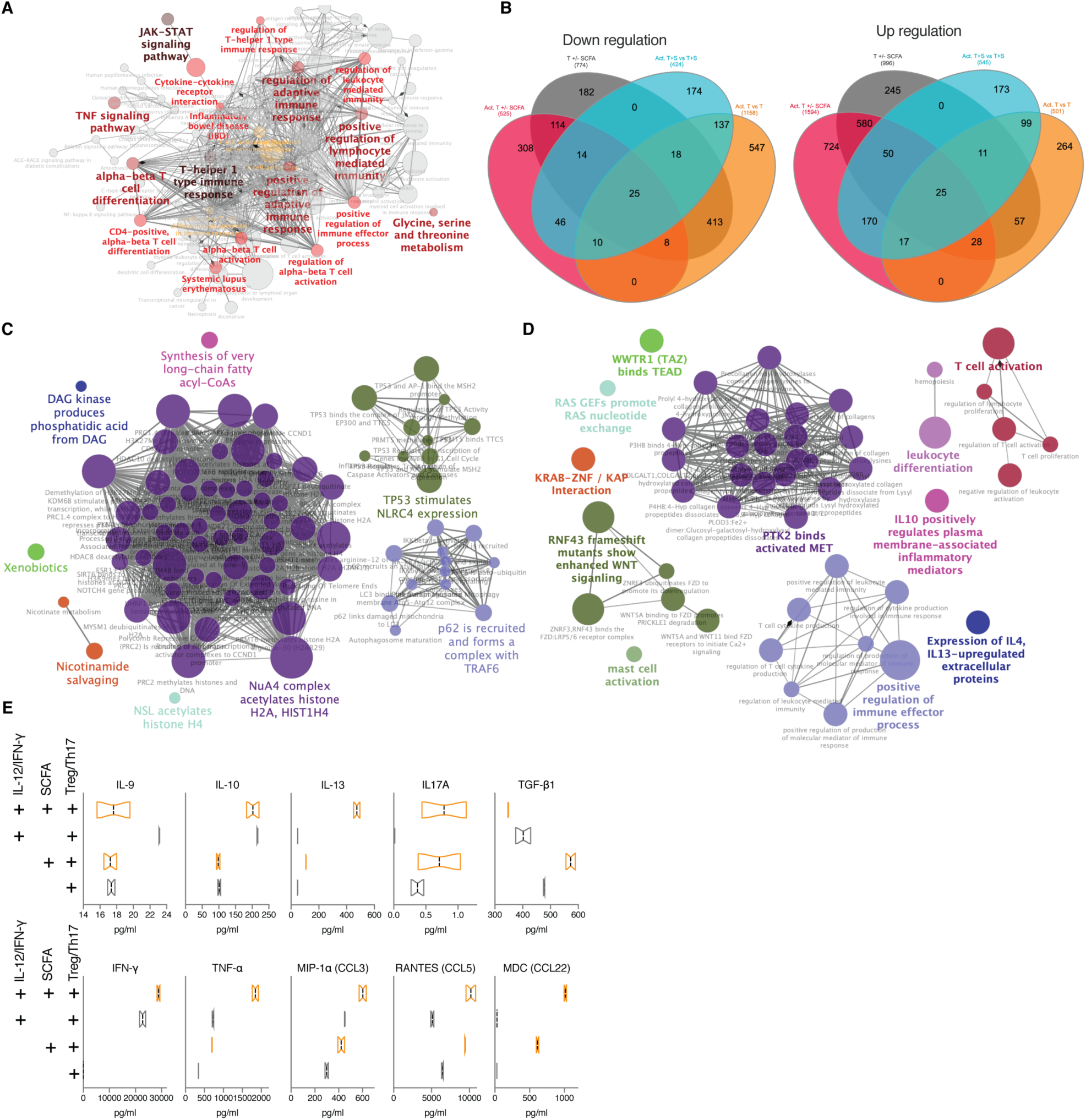
SCFA promote production of IL-13 and other inflammatory cytokines exclusively in activated T cells. (A) Network of significantly enriched pathways based on KEGG in Treg/Th17 cells stimulated with IL-12 and IFN-γ (B) Venn diagram showing shared and unique significantly downregulated (left) or upregulated (right) genes among IL-12/IFN-γ stimulated Treg/Th17 cells +/- 1mM of total SCFA (magenta), unstimulated Treg/Th17 cells +/- 1mM of total SCFA (grey), IL-12/IFN-γ stimulated Treg/Th17 cells with SCFA over unstimulated T cells with SCFA (cyan), IL-12/IFN- γ stimulated Treg/Th17 cells over unstimulated T cells (orange). (C,D) Representative networks of downregulated pathways (C) or upregulated pathways (D) enriched based on REACTOME exclusively in IL-12/IFN-γ stimulated Treg/Th17 cells treated with 1mM of total SCFA. RNAseq analysis was performed on pooled cells from 3 independent experiments. (E) Multiplexed cytokines/chemokines in supernatants collected from each stimulated or unstimulated group with or without SCFA after 48 h of incubation.

Enrichment based on the REACTOME database indicated strong downregulation of HDAC activity, which has been shown in multiple studies (Furusawa et al., 2013), and p62-TRAF6 autophagy related pathways in activated, SCFA-treated CD4 T cells exclusively (Fig.7C). TRAF6 is an intrinsic T cell regulator required for immune homeostasis and its deletion leads to multiorgan inflammatory disease (King et al., 2006). Simultaneously, WNT activity and ECM receptor activity were increased in the same group (Fig.7D). T cell activation pathways were significantly increased among stimulated and SCFA treated CD4 T cells, which is in line with a recent study on murine T cells (Kim et al., 2013). Moreover, SCFA were shown to inhibit FOXP3, the developmental gene required for Treg differentiation, thus dictating CD4 T cell effector versus tolerance phenotypes depending on the activation level of the cell (Park et al., 2015). Specifically, both IL-10 and IL-13 cytokine-regulated extracellular protein pathways were increased as also seen during the interaction study with Treg/Th17 cells (Fig.S6).

Cytokine/chemokine measurements of their supernatants revealed stark differences among the groups (Fig.7E). Under steady state conditions addition of SCFA led to strongly increased TGF-β1 production consistent with previously published observations, in fact promoting tolerance. However, the trend seemed to be reversed in activated T cells, where SCFA led to increased release of IFN-γ, TNF-α and in particular IL-13, confirming the exacerbation potential of SCFA under inflammatory conditions. IL-13 is a critical promoter of UC and a target of interest in therapies against UC (Hoving, 2018). Interestingly, regardless of activation state, SCFA prompted the higher release of the chemokines RANTES and MDC, which are important for additional recruitment of lymphocytes to sites of infection. Identifying the ability of SCFA to significantly increase IL-13 production by activated T cells and inhibit p62-TRAF6 activity is a novel finding with broad implication in understanding UC.

## Discussion

By including Treg/Th17 cells in the interaction between the UC gut MPSs and liver MPSs, we recapitulated acute inflammatory conditions and global metabolomic changes described in advanced UC with concurrent autoimmune hepatic disease. While the mismatch between donors of individual cell types used in the study represents a technical limitation, we do believe the system to accurately represent the human biological CD4 T cell response, due to known disease parallels between alloimmunity, IBD or autoimmune liver diseases (Belyayev et al., 2019; Carbone and Neuberger, 2014). Hence, the impact of the mismatch on the observed outcomes is likely minimal. The gut MPS-induced CD4 T cell-mediated acute inflammation allowed us to probe temporal interconnectedness of microbiome-derived SCFA, innate and adoptive immune response without the use of xenopeptides to induce autoreactivity or chemicals, routinely used in animal models of IBD and autoimmune hepatitis (Erben et al., 2014; Liu et al., 2013; Taneja and David, 2001). However, further advances in hiPSC and other technologies will allow us to recreate donor-matched physiomimetic experiments for broader spectrum of diseases in the future.

In partial agreement with previously published data, we were able to confirm SCFA to exert wide-ranging modifications of human gut and liver function, in part through actions on the innate and adaptive immune systems. SCFA appear to decrease innate immune activation of the ulcerative colitis gut MPS through PPAR signaling and downregulation of the NF-kappa B pathway (Fig.1-4). Reduced inflammation of the gut MPS favorably affected the gut-liver axis with increased hepatic metabolic function where conversion of SCFA lead to increased bile acid production, gluconeogenesis, lipid metabolism and formation of ketone bodies (Fig.4). In contrast to this observation and common belief, SCFA gravely exacerbated CD4 T effector cell activation and liver injury (Fig.5-7). This was largely dependent on the direct influence of SCFA on activated Treg/Th17 cells through HDAC and p62-TRAF6 inhibition, metabolic reprograming and increased differentiation towards Th1/Th2 effector cells. At the same time, SCFA did promote Treg differentiation and production of TGF-β in non-stimulated Treg/Th17 populations as previously reported. Moreover, we have discovered the ability of SCFA to strongly increase the production of IL-13, a cytokine tightly connected to severity and progression of UC in humans. Cumulatively, these findings indicate that the role of SCFA greatly depends on the activation state of receptive CD4 T cells. While SCFA reduce innate immune activation of primary tissue in the absence of acute T cell inflammation, such as during early stages of UC, they might exert opposing effects in progressed acute UC with negative consequences for the hepatobiliary system. Activated, highly proliferative T cells largely require glucose for a robust response while resting immune cells switch fuel usage from glucose to fatty acids and ketones in support of tissue-protective pathways (Wang et al., 2019). However, SCFA and increased production of ketone bodies measured during interaction studies, further increased T cell effector function. This indicates that internal lymphocyte metabolic reprograming might not be as black and white, and that both anabolic and catabolic metabolic programs are potentially engaged concurrently which should be further explored. Our findings might explain in part the plethora of conflicting reports on the effects of SCFA in autoimmune/autoinflammatory diseases where their action seems to be highly context dependent.

Recreating hallmarks of human innate and adaptive immune mechanics of the gut-liver axis, although in absence of gut microbes, highlights the potential of human physiomimetic technologies to fill in the gaps in studying complex diseases through mechanistic correlation of various multiomic observations and systems immunology to both human and animal clinical data.

## Acknowledgments

The authors are grateful to D Lauffenburger, S Mazmanian, A Wells for their critical input; to B Ringeisen, D Stepp, R Cecil and G Kost for their support and feedback; D Breault and F Zhou of the Harvard Digestive Disease Center as well as J Papps and V Hernandez-Gordillo for their help with establishing intestinal organoid cultures; V Butty, S Levine, D Brubaker and J Das for their help with RNAseq analysis and data representation; to JW Kemmitt for his help in operating the physiomimetic platforms and H Lee for managing lab operations

## Funding

The study was funded by the grants DARPA W911NF-12-2-0039, NIH/NIBIB R01EB021908 and in part by the National Institute of Environmental Health Sciences of the NIH under award P30-ES002109, the NIH grant P30DK034854 as well as NIH R01 CA211184, CA034992 and the Pew-Stewart Trust Foundation.

## Data and materials availability

Raw data and materials can be obtained by inquiry to the authors.

## Competing interests

Authors declare no competing interests;

## Author contributions

MT, CC, OY, DT and LG were responsible for conceptualization, investigation, data curation and analysis, methodology, visualization, and writing and review. MT, JV, CAM, YH, KS, CWW and GE performed the experiments and assisted with data analysis.

## STAR Methods

### Contact for Reagent and Resource Sharing

Information and requests for resources and reagents should be directed to and will be fulfilled by the Lead Contact, Linda G. Griffith

### Experimental Model and Subject Details

#### Gut microphysiological systems (gut MPS)

##### Gut organoids

Colon organoids used in this study were established and maintained as previously described (Kasendra et al., 2018; Roper et al., 2017) by the Harvard digestive disease center (healthy control donor – 6 months old female) and the Yilmaz lab at the Koch Institute/MIT (UC donor – sex and age not disclosed). Endoscopic tissue biopsies were collected from the ascending colon of de-identified individuals at either Boston Children’s Hospital or Massachusetts General Hospital upon the donors informed consent. Methods were carried out in accordance to the Institutional Review Board of Boston Children’s Hospital (protocol number IRB-P00000529) and the Koch Institute Institutional Review Board Committee as well as the Massachusetts Institute of Technology Committee on the Use of Humans as Experimental Subjects. Tissue was digested in 2 mg ml^−1^ collagenase I (StemCell, cat. no. 07416) for 40 min at 37 °C followed by mechanical dissociation, and isolated crypts were resuspended in growth factor-reduced Matrigel (Becton Dickinson, cat. no. 356237) and polymerized at 37 °C. Organoids were grown in expansion medium (EM) consisting of Advanced DMEM/F12 supplemented with L-WRN conditioned medium (65% vol/vol, ATCC, cat. no. CRL-3276), 2 mM Glutamax (Thermo Fisher, cat. no. 35050-061), 10 mM HEPES (Thermo Fisher, cat. no. 15630-080), Penicillin/Streptomycin (Pen/Strep) (Thermo Fisher, cat. no. 15070063), 50 ng ml^−1^ murine EGF (Thermo Fisher, cat. no. PMG8041), N2 supplement (Thermo Fisher, cat. no. 17502-048), B-27 Supplement (Thermo Fisher, cat. no. 17502-044), 1 nM human [Leu15]-gastrin I (Sigma, cat. no. G9145), 0.5 mM N-acetyl cysteine (Sigma, cat. no. A9165-5G), 10 mM nicotinamide (Sigma, cat. no. N0636), 2.5 µM thiazovivin (Tocris, cat. no. 3845), 10 µM SB202190 (Tocris, cat. no. 1264), 500 nM A83-01 (Tocris, cat. no. 2939), 5 nM prostaglandin E2 (StemCell cat. no. 72192) at 37°C and 5% CO_2_.

Organoids were passaged every 7 days by incubating in Cell Recovery Solution (Corning, cat. no. 354253) for 40 min at 4 °C, followed by mechanical dissociation and reconstitution in fresh Matrigel at a 1:4 ratio.

##### Epithelial monolayers on Transwell inserts

At day 7 post passaging, colon organoids were collected, Matrigel was dissolved with Cell Recovery Solution for 40 min at 4 °C followed by incubation of Matrigel-free organoids in TrypLE Express (Thermo Fisher, cat. no. 12605036) at 37 °C for 5 minutes. Organoids of both donors were mechanically dissociated into single cells, resuspended in EM without nicotinamide and seeded onto type I collagen/Matrigel coated 12-well 0.4 µm pore polyester Transwell inserts (Corning, 3493) at a density of 3 x 10^5^ cells/Transwell. After 3-4 days of incubation, monolayers were confluent and differentiation was initiated. For differentiation apical media was replaced with Advanced DMEM/F12 plus Glutamax, HEPES and Pen/Strep and basalolateral media with differentiation medium (DM), which is EM without L-WRN conditioned medium, nicotinamide, prostaglandin E2 and thiazovivin, but supplemented with 100 ng ml^−1^ human recombinant noggin (Peprotech, cat. no. 120-10C) and 20% R-spondin conditioned media (Sigma, cat. no. SCC111). Transepithelial electrical resistance (TEER) measurements were performed using the EndOhm-12 chamber with an EVOM2 meter (World Precision Instruments). At day 8 post seeding, the monolayers were used for further experimentation.

##### Coculture of epithelial monolayers with dendritic cells and macrophages

Human monocyte-derived dendritic cells and macrophages were used as the innate immune component of the gut MPS and seeded on the basolateral side of Transwell membranes that have epithelium on the apical side. Monocytes were isolated from peripheral blood mononuclear cells (PBMCs) containing Leuko Pak of a 44 years old Caucasian female (StemCell, cat. no. 70500) using the EasySep Human Monocyte Enrichment Kit (StemCell, cat. no. 19058). Dendritic cells were differentiated in RPMI 1640 medium (Gibco) supplemented with Pen/Strep, heat-inactivated 10% heat-inactivated FBS, 1% MEM Non-Essential Aamino Acid Solution (Gibco), 1% GlutaMax, 100 ng/mL GM-CSF (Biolegend, cat. no. 572903), 70 ng/mL IL-4 (Biolegend, cat. no. 574004) and 10 nM retinoic acid (Sigma, cat. no. R2625-50MG). Macrophages were differentiated in RPMI 1640 medium (Gibco) supplemented with 10% FBS, 1x GlutaMax, 100 ng/mL M-CSF (Biolegend, cat. no. 716602). After 7 days of differentiation (at day 8 post epithelial cell seeding), dendritic cells and macrophages were harvested using Accutase (Gibco) and seeded onto the basal side of the Transwells, 2.5×10^4^ of each population per Transwell. Gut MPS with TEER greater than 200 ohm/cm^2^ were considered acceptable for experimentation and were integrated onto the 3XGL platform for interaction studies or studies in isolation. During all experiments in isolation or interaction, the gut MPS was maintained in serum-free apical medium consisting of Advanced DMEM/F12 with or without the short-chain fatty acids (SCFA), sodium acetate (12mM), sodium propionate (4mM) and sodium butyrate (4mM) from Sigma-Aldrich. The basal gut compartment as well as the liver and mixer compartment that were fluidically linked to systemic circulation on the 3XGL were fed with serum-free common media (CM) that contained William’s E medium (Thermo Fisher, cat. no. A1217601), 4% Cell Maintenance Supplement Pack (Thermo Fisher, cat. no. CM4000), 50 UI/ml IL-2 (R&D Systems, cat. no. 202-IL), 100 nM hydrocortisone, 5 mM glucose, 800 pM insulin and 0.5% Pen/Strep. Experiments here and throughout the entire work were conducted in a slight modification of culture medium which had previously been tailored for physiological inflammatory responses of the human liver MPS, in which the extreme supra-physiological concentrations of cortisol and insulin typically used to maintain CYP450 levels in primary hepatocyte cultures were reduced to concentrations within the physiological range (Long et al., 2016; Sarkar et al., 2015).

#### Liver microphysiological systems (liver MPS)

The liver MPS was prepared as previously described (Chen et al., 2017). Single donor human primary hepatocytes were purchased from BioIVT (lot AQL; 63-years old male) and Kupffer cells, as the innate immune component of the liver MPS, were obtained from Thermo Fisher (cat. no. HUKCCS; 29-years old male). The cells were seeded on liver scaffolds which are thin (0.25 mm) polystyrene discs perforated with 301 channels (diameter = 0.3 mm) (Clark et al., 2016). Scaffolds were soaked in 70% EtOH for 15 min, washed twice with PBS, and coated with 30 μg/mL rat tail collagen I in PBS for 1 hour at room temperature. Collagen-coated scaffolds were air dried and then punched into the 3XGL platforms for interaction experiments or in the LiverChip® (CN Bio Innovations) for experiments in isolation. 3 days prior to interaction experiments, hepatocytes and Kupffer cells were thawed in Cryopreserved Hepatocyte Recovery Medium (Thermo Fisher, cat. no. CM7000), centrifuged at 100 × g for 8 min, and seeded on the scaffolds at a 10:1 ratio, 6 × 10^5^:6 × 10^4^ cells/well, in hepatocyte seeding medium (William’s E medium (Thermo Fisher, cat. no. A1217601) with 5mM Glucose, 5% FBS, 100nM hydrocortisone, and an in house supplement cocktail equivalent to Gibco Cocktail A (Thermo Fisher, cat. no. CM3000) but with only 200-800pM insulin) and cultured under flow at 37°C. After 1 day, the media was changed to hepatocyte maintenance medium (William’s E medium with 5mM glucose, 100nM hydrocortisone, and an in house supplement cocktail equivalent to Gibco Cocktail B (Thermo Fisher, cat. no. CM4000) but with only 200-800pM insulin). At day 3 after seeding, the media was changed to CM + 25UI IL-2 and the interaction studies with the gut MPS begun. The CM was also added to the LiverChip®. To evaluate the physiological status of the liver, samples from all compartments but the apical gut MPS (i.e., liver, directly above the scaffold; basal gut MPS: and mixer) were taken at every medium change (every 48 hours) and assayed for albumin via ELISA (Bethyl Laboratories, cat. no. E80-129).

#### Human circulating CD4 regulatory T cells (Treg) and Th17 cells

Treg and Th17 cells were differentiated from naïve CD4 T cells derived from the same Leuko Pak donor, a 44 years old Caucasian female (StemCell, cat. no. 70500), as the dendritic cells and macrophages in the gut MPS. Naïve CD4 T cells were isolated with the EasySep Human Naïve CD4+ T Cell Isolation Kit II (StemCell, cat. no. 17555). Cells were differentiated to Tregs or Th17 cells in RPMI 1640 medium (Gibco) supplemented with Pen/Strep, heat-inactivated 10% heat-inactivated FBS, 1% MEM Non-Essential Amino Acid Solution, 1 mM sodium pyruvate, 1% GlutaMax, 2.5% ImmunoCult Human CD3/CD28 T cell activator (StemCell, cat. no. 10971) and 5 ng/ml human TGF-beta (R&D Systems, cat. no. 240-B/CF). Additionally Tregs received 100 UI/ml IL-2 (R&D Systems, cat. no. 202-IL) and 10 nM retinoic acid (Sigma, cat. no. R2625-50MG); while for Th17 cell differentiation, 10 ng/ml each of human IL-6 and IL-1 beta (R&D Systems, cat. no. 206-IL, 201-LB) were added. After 7 days of differentiation at 37 °C, cells were harvested into CM and added in a physiological 2:1 ratio of Treg/Th17 cells to either the basal side of gut MPS for isolation studies (total of 4 x 10^4^ Tregs/2 x 10^4^ Th17 cells/Transwell) or equally distributed among the compartments on the 3XGL platform (total of 2.4 x 10^5^ Tregs and 1.2 x 10^5^ Th17 cells/interaction lane). Treg/Th17 ratio in human peripheral blood varies from 0.2 to 1 in healthy individuals and total Treg numbers varying from 7 x 10^4^ to 5 x 10^5^/ml (Eastaff-Leung et al., 2010; Ratajczak et al., 2010). While T cells were prevented to come into direct contact with the epithelium grown on microporous membranes, their contact was enabled with antigen presenting cells of the gut MPS and the liver MPS.

#### Physiomimetic platform 3XGL

##### Fabrication and assembly

The physiomimetic technology was developed in-house and described previously (Chen et al., 2017; Edington et al., 2018). Briefly, iterations of the system were designed in CAD and commercially machined. The pneumatic plates were machined in acrylic and solvent bonded to form two-layer manifolds, while the fluidic plates were machined from monolithic polysulfone. The pneumatic plates were processed with vapor polishing and fluidic plates were cryo-deburred to remove sharp burrs. 50 micron thick polyurethane membranes are supplied by American Polyfilm Inc. and mounted onto grip rings (Ultron Systems UGR-12) to provide uniform tension. Mounted membranes were laser cut to remove material around screw holes. After cutting, they were rinsed in 10% 7x solution followed by DI water and then sterilized using ethylene oxide gas. All hardware components, with the exception of the polyurethane membranes, are reusable. Flow of media between compartments on the platform is achieved by pumping driven from outside the incubator by a microcontroller and a pneumatic solenoid manifold that controls the tubing run through the back of the incubator to intermediary connectors. Moreover, the system allows for safe circulation of CD4 T cells within and between compartments at high velocities and preserved T cell viability. Inside the incubator, tubing is attached to the platform through valved breakaway couplings to allow removal from the incubator for media changes and sampling. Flow rates and calibration factors are set through a graphical user interface and are sent to a customized microcontroller (National Instruments, cat. no. myRIO-1900) over USB or WiFi.

### Method Details

#### Operation of the 3XGL physiomimetic platform and MPS interaction studies

4 days prior to experimentation, sterile platforms were assembled in a laminar flow hood. They were primed with 1% bovine serum albumin (Sigma, cat. no. A9576) and Pen/Strep in PBS. Pump function and tubing connections were visually confirmed by pumping from the mixer to each dry compartment, then by running the recirculation pumps backwards to clear all air from the channels. Platforms were run overnight in the incubator to passivate and confirm full operation before the addition of the gut and/or liver MPS as well as the circulating Treg/Th17 cells. On the day of the experiment, priming media was replaced with serum-free CM (see Gut MPS methods) with the volume of each compartment being as follows: gut MPS 2 ml (apical 0.5 ml, basal 1.5 ml), liver MPS 1.6 ml and the mixer compartment 2.3 ml. The layout of the platform and flow parameters are indicated in Figure S2A. The duration of interaction studies was 4 days with complete media changes every 48 hours. TEER was measured just prior to the interaction and at the end on day 4 to minimize monolayer manipulation. However, TEER was measured on control gut MPSs in isolation at day 2 to monitor innate monolayer behavior. During the media change and at the end of the interaction studies on day 4, common media was collected from the apical and basal gut compartment, liver compartment above the scaffold and mixer. Circulating Treg/Th17 cells that were collected with the media during the 48 h media change were returned to each original platform equally distributed among the 3 compartments. Media samples were collected in low-binding tubes, supernatants were spun down at 10,000 g for 5 min to remove cell debris, BSA was added to a final concentration of 0.5% (except to the samples reserved for metabolomic analysis) and the samples transferred to a -80 °C freezer. Cytokine/chemokine, metabolomic and albumin measurements during interaction studies were performed on the media collected from the apical gut compartments (for studies of SCFA distribution only) and the mixer compartments that distribute the media between the gut and liver MPSs. Each condition was performed in 3 biological replicates.

#### Multiplex cytokine/chemokine assays

Cytokine levels were measured using the following multiplex cytokine assays from Millipore Sigma: MILLIPLEX MAP Human Cytokine/Chemokine Magnetic Bead Panel - Premixed 41 Plex (cat. no. HCYTMAG-60K-PX41), MILLIPLEX MAP TGFß Magnetic Bead 3 Plex Kit (cat. no. TGFBMAG-64K-03) and a custom MILLIPLEX MAP Human Th17 Panel (cat. no. HTH17MAG-14K-10). We reconstituted the protein standard in the same media and serially diluted the protein stock to generate an 8-point standard curve. A Bio-Plex 3D Suspension Array System (Bio-Rad Laboratories, Inc.) was used to analyze the samples.

#### Treg and Th17 cell activation assay

To investigate the effect of SCFA on the function of Treg and Th17 cells, differentiated cells were seeded in combination (2.4×10^5^ cells/ml of Tregs and 1.2×10^5^ cells/ml of Th17 cells) in 1ml of CM on 24-well plates. Cells were challenged with either 1mM of total SCFA (0.7mM acetate, 0.2mM propionate, 0.1mM butyrate); 10 ng/ml of IL-12 and IFN-γ (R&D Systems, cat. no 212-IL/285-IF); or the combination of SCFA, IL-12 and IFN-γ. Cells were treated for 48 hours at 37°C in 5% CO_2_. Supernatants were stored for multiplex cytokine analysis and cells were harvested for RNAseq analysis.

#### Immunofluorescence confocal imaging

Transwell membranes containing epithelial monolayers on the apical side and macrophages/DCs on the basolateral side were washed with FACS buffer (PBS with 2%FBS) and stained with primary mouse anti-human CD14 Clone MφP9 (BD Biosciences, cat. no. 347490). After washing with FACS buffer cells were fixed and permeabilized with a Fixation/Permeabilization kit (BD Biosciences, cat. no. 554714) and concurrently stained with an AlexaFluor 488 secondary goat anti-mouse antibody (Biolegend, cat. no. 405319), NucBlue (Invitrogen, cat. no. 12303553) and ActinRed (Invitrogen, cat. no. 15119325). Membranes were mounted on glass slides with Prolong Diamond Antifade (Invitrogen, cat. no. 15205739) and cured for 24 hours. Images were acquired with the Zeiss LSM 880 confocal microscope at a 63x magnification and curated with the Zeiss ZEN software.

#### Metabolomics

Targeted SCFA and broad discovery metabolomic analysis as well as bioinformatic data processing were performed by Metabolon (Metabolon Inc., Durham, NC, United States) while PBPK modeling of SCFA was performed in-house. Samples from the experiments were subject to metabolite extractions and analysis by RP/UPLC-MS/MS (ESI+) (-ESI) with details of the methods published previously (Evans AM, 2014). Following receipt, samples were inventoried and immediately stored at -80°C. Each sample received was accessioned into the Metabolon LIMS system and assigned by the LIMS a unique identifier that was associated with the original source identifier only. This identifier was used to track all sample handling, tasks, results, etc. The samples (and all derived aliquots) were tracked by the LIMS system. All portions of any sample were automatically assigned their own unique identifiers by the LIMS when a new task was created; the relationship of these samples was also tracked. All samples were maintained at -80°C until processing.

##### Determining concentrations of SCFA

Media samples are analyzed for eight short chain fatty acids: acetic acid (C2), propionic acid (C3), isobutyric acid (C4), butyric acid (C4), 2-methyl-butyric acid (C5), isovaleric acid (C5), valeric acid (C5) and caproic acid (hexanoic acid, C6) by LC-MS/MS (Metabolon Method TAM131). Media samples were spiked with stable labelled internal standards and are subjected to protein precipitation with methanol. After centrifugation, an aliquot of the supernatant was derivatized. The reaction mixture was diluted, and an aliquot was injected into an Agilent 1290/AB Sciex QTrap 5500 LC MS/MS system equipped with a C18 reversed phase UHPLC column. The mass spectrometer was operated in negative mode using electrospray ionization (ESI). The peak area of the individual analyte product ions was measured against the peak area of the product ions of the corresponding internal standards. Quantitation was performed using a weighted linear least squares regression analysis generated from fortified calibration standards prepared immediately prior to each run. Solutions and calibration standards were prepared per Method TAM131. Single level pooled media QC samples were used with all analytes at the endogenous levels. Due to the demands of multiplexing eight analytes, only a single concentration level of QC samples was used.

Raw data were collected and processed using AB SCIEX software Analyst 1.6.2. Data reduction was performed using Microsoft Excel 2013. Sample analysis was carried out in the 96-well plate format containing two calibration curves and six QC samples (per plate) to monitor method performance.

Precision was evaluated using the corresponding QC replicates in each sample run. Intra-run precision (%CV) of all analytes met acceptance criteria. (Targeted acceptance criteria for intra-run QC precision: %CV ≤ 20%.).

##### Global metabolomic discovery

We have performed the global metabolomic discovery on control common media, media collected from the mixer compartments during interaction studies and media collected from control gut and liver MPSs in isolation as described under “Operation”. Samples were prepared using the automated MicroLab STAR® system from Hamilton Company. Several recovery standards were added prior to the first step in the extraction process for QC purposes. To remove protein, dissociate small molecules bound to protein or trapped in the precipitated protein matrix, and to recover chemically diverse metabolites, proteins were precipitated with methanol under vigorous shaking for 2 min (Glen Mills GenoGrinder 2000) followed by centrifugation. The resulting extract was divided into five fractions: two for analysis by two separate reverse phase (RP)/UPLC-MS/MS methods with positive ion mode electrospray ionization (ESI), one for analysis by RP/UPLC-MS/MS with negative ion mode ESI, one for analysis by HILIC/UPLC-MS/MS with negative ion mode ESI, and one sample was reserved for backup. Samples were placed briefly on a TurboVap® (Zymark) to remove the organic solvent. The sample extracts were stored overnight under nitrogen before preparation for analysis.

Several types of controls were analyzed in concert with the experimental samples: a pooled matrix sample generated by taking a small volume of each experimental sample (or alternatively, use of a pool of well-characterized human plasma) served as a technical replicate throughout the data set; extracted water samples served as process blanks; and a cocktail of QC standards that were carefully chosen not to interfere with the measurement of endogenous compounds were spiked into every analyzed sample, allowing instrument performance monitoring and aiding chromatographic alignment. Instrument variability was determined by calculating the median relative standard deviation (RSD) for the standards that were added to each sample prior to injection into the mass spectrometers. Overall process variability was determined by calculating the median RSD for all endogenous metabolites (i.e., non-instrument standards) present in 100% of the pooled matrix samples. Experimental samples were randomized across the platform run with QC samples spaced evenly among the injections.

All methods utilized a Waters ACQUITY ultra-performance liquid chromatography (UPLC) and a Thermo Scientific Q-Exactive high resolution/accurate mass spectrometer interfaced with a heated electrospray ionization (HESI-II) source and Orbitrap mass analyzer operated at 35,000 mass resolution. The sample extract was dried then reconstituted in solvents compatible with each of the four methods. Each reconstitution solvent contained a series of standards at fixed concentrations to ensure injection and chromatographic consistency. One aliquot was analyzed using acidic positive ion conditions, chromatographically optimized for more hydrophilic compounds. In this method, the extract was gradient eluted from a C18 column (Waters UPLC BEH C18-2.1×100 mm, 1.7 µm) using water and methanol, containing 0.05% perfluoropentanoic acid (PFPA) and 0.1% formic acid (FA). Another aliquot was also analyzed using acidic positive ion conditions, however it was chromatographically optimized for more hydrophobic compounds. In this method, the extract was gradient eluted from the same afore mentioned C18 column using methanol, acetonitrile, water, 0.05% PFPA and 0.01% FA and was operated at an overall higher organic content. Another aliquot was analyzed using basic negative ion optimized conditions using a separate dedicated C18 column. The basic extracts were gradient eluted from the column using methanol and water, however with 6.5mM ammonium bicarbonate at pH 8. The fourth aliquot was analyzed via negative ionization following elution from a HILIC column (Waters UPLC BEH Amide 2.1×150 mm, 1.7 µm) using a gradient consisting of water and acetonitrile with 10mM ammonium formate, pH 10.8. The MS analysis alternated between MS and data-dependent MSn scans using dynamic exclusion. The scan range varied slighted between methods but covered 70-1000 m/z. Raw data files are archived.

#### RNAseq

Library preparation, sequencing and analysis was performed by the BioMicro Center at MIT.

##### RNA extraction, cDNA library preparation and next generation sequencing

Epithelial tissue from the gut MPSs and hepatocytes with Kupffer cells from the liver MPSs, as well as Treg/Th17 cells were collected and mRNA was extracted using the PureLink RNA mini kit (Thermo Fisher, cat. no. 12183018A). Total RNA was analyzed and quantified using the Fragment Analyzer (Advanced Analytical), and Illumina libraries were prepared with Kapa RNA HyperPrep with RiboErase (Kapa Biosystems, cat. no. 08098107702). After cDNA fragmentation (Covaris S2), cDNA was end-repaired and adaptor-ligated using the SPRI-works Fragment Library System I (Beckman Coulter Genomics). Adaptor-ligated cDNA was then indexed during PCR amplification, and the resulting libraries were quantified using the Fragment Analyzer and qPCR before being sequenced on the Illumina HiSeq 2000 (Illumina). 4 lanes 40-50 nt single-end read with an average depth of 15-20 million or 5 million reads per sample were sequenced.

### Quantification and Statistical Analysis

#### Analysis of multiplexed cytokine/chemokine data

Data from the multiplexed cytokine/chemokines was collected using the xPONENT for FLEXMAP 3D software, version 4.2 (Luminex Corporation, Austin, TX, USA). The concentration of each analyte was determined from a standard curve, which was generated by fitting a 5-parameter logistic regression of mean fluorescence on known concentrations of each analyte (Bio-Plex Manager software). Hierarchical complete clustering, heatmaps and principal component analysis of cytokine MFIs were performed using ClustVis (Metsalu and Vilo, 2015), an online platform integrating several R packages for analysis. Cytokine data were normalized by mean-centering and variance scaling prior to clustering and principal component analysis. Actual cytokine concentrations were plotted in Prism 8.0 (GraphPad Software) as well as their log2 fold change comparing individual experimental groups. Unpaired *t* test was used to calculate statistical significance. Moreover, the importance of variables was determined by the machine learning algorithm random forest using the software BioVinci (BioTuring Inc.).

#### Bioinformatic analysis of identified metabolomic targets

The Metabolon informatics system consisted of four major components, the Laboratory Information Management System (LIMS), the data extraction and peak-identification software, data processing tools for QC and compound identification, and a collection of information interpretation and visualization tools for use by data analysts. The hardware and software foundations for these informatics components were the LAN backbone and a database server running Oracle 10.2.0.1 Enterprise Edition.

Raw data were extracted, peak-identified and QC processed using Metabolon’s hardware and software. These systems are built on a web-service platform utilizing Microsoft’s .NET technologies, which run on high-performance application servers and fiber-channel storage arrays in clusters to provide active failover and load-balancing. Compounds were identified by comparison to library entries of purified standards or recurrent unknown entities. Metabolon maintains a library based on authenticated standards that contains the retention time/index (RI), mass to charge ratio (m/z), and chromatographic data (including MS/MS spectral data) on all molecules present in the library. Furthermore, biochemical identifications are based on three criteria: retention index within a narrow RI window of the proposed identification, accurate mass match to the library +/- 10 ppm, and the MS/MS forward and reverse scores between the experimental data and authentic standards. The MS/MS scores are based on a comparison of the ions present in the experimental spectrum to the ions present in the library spectrum. While there may be similarities between these molecules based on one of these factors, the use of all three data points can be utilized to distinguish and differentiate biochemicals. More than 3300 commercially available purified standard compounds have been acquired and registered into LIMS for analysis on all platforms for determination of their analytical characteristics. Additional mass spectral entries have been created for structurally unnamed biochemicals that have been identified by virtue of their recurrent nature (both chromatographic and mass spectral). These compounds have the potential to be identified by future acquisition of a matching purified standard or by classical structural analysis.

Peaks were quantified using area-under-the-curve. For samples analyzed on different days, a data normalization step was performed to correct variation resulting from instrument inter-day tuning differences. Essentially, each compound was corrected in run-day blocks by registering the medians to equal one (1.00) and normalizing each data point proportionately (termed the “block correction”). For studies that did not require more than one day of analysis, no normalization is necessary, other than for purposes of data visualization. In certain instances, biochemical data may have been normalized to an additional factor (e.g., cell counts, total protein as determined by Bradford assay, osmolality, etc.) to account for differences in metabolite levels due to differences in the amount of material present in each sample. Two types of statistical analysis are usually performed: (1) significance tests and (2) classification analysis. Standard statistical analyses are performed in ArrayStudio on log transformed data. For those analyses not standard in ArrayStudio, the programs R (http://cran.r-project.org/) or JMP were used.

Pathway enrichment analysis and visualization was performed with Metabolons proprietary Pathway Explorer tool.

#### PBPK Models of intestinal SCFA bioaccessibility and biodistribution through the 3XGL platform

Intestinal clearance and bioaccessibility of SCFA was investigated on the 3XGL platforms without the liver MPS or circulating Treg/Th17 cells under self-circulation rates identical to the operation during the interaction study. Media from the apical and basal sides of the gut MPS were collected at 4h, 8h, 24h and 48h after application of 20 mM total SCFA to the apical compartment at a physiological ratio of 6:2:2 (acetate:propionate:butyrate). A mathematical model was used to determine SCFA distribution, metabolism, and permeability for all gut MPS. A two-compartmental model was used to describe the apical and basal site of the gut MPS (V_apical_ = 0.5 ml, V_basal_ = 1.5 ml) (Fig.S2B). The model was used to simultaneously describe both apical and basal measured SCFA concentrations (acetate, propionate, butyrate) over the course of the 48h experiment by fitting permeability of SCFA between the compartments (*P_gut_*) as well as metabolism of individual SCFA from the apical compartment (*CL*).

Compartent #1: apical

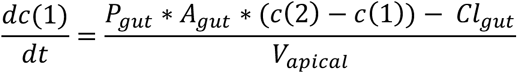

Compartment #2: basal

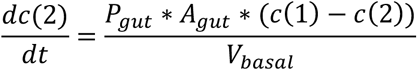

Further distribution and liver MPS clearance of SCFA during gut-liver-immune interaction was determined with a four-compartmental model. In addition to the aforementioned estimated gut permeability and metabolism, metabolism of SCFA by the liver MPS was implemented (*CL_L_*) (Fig.S2C).

All parameters were estimated by minimizing the squared difference (least-square) between actual individual SCFA measurements and the model description for compartments and SCFAs simultaneously.

**Table S1.** Overview of model parameters

| Parameter name | Acronym | Value | Unit |
| --- | --- | --- | --- |
| Permeability | $P_{gut}$ | Estimated | cm/min |
| Surface area | $A_{gut}$ | 0.33 | cm<sup>2</sup> |
| Metabolism gut | $Cl_{gut}$ | Estimated | ml/min |
| Metabolism liver | $Cl_{liv}$ | Estimated | ml/min |
| Media volume gut | $V_{apical}$ | 0.5 | ml |
| | $V_{basal}$ | 1.5 | ml |
| Media volume liver | $V_{liv}$ | 1.6 | ml |
| Media volume mixer | $V_{mix}$ | 2.5 | ml |
| Systemic flow rate | $Q_{mix}$ | 30 | ml/day |
| Flow partitioning gut | $Q_{gut}$ | $0.75 * Q_{mix}$ | ml/day |
| Flow partitioning liver | $Q_{hep}$ | $0.25 * Q_{mix}$ | ml/day |

The resulting parameter values were averaged over three biological replicates and the corresponding standard deviation was calculated.

Compartment #1: mixing chamber

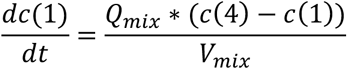

Compartment #2: gut apical

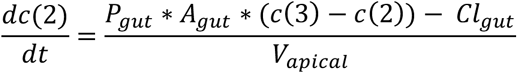

Compartment #3: gut basal

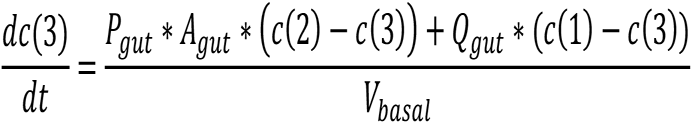

Compartment #4: liver

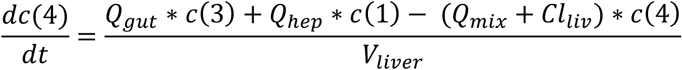

Model performance was assessed by visual inspection of simulated and measured SCFA time-concentration profiles. All simulations and parameter estimations were performed in MATLAB (R2018a, The MathWorks, Inc., Natick, Massachusetts, United States).

#### RNAseq data analysis

Quality control: Reads were aligned against the hg19 (Feb 2009) human genome assembly using bwa mem v. 0.7.12-r1039 [http://bio-bwa.sourceforge.net/] with flags –t 16 –f and mapping rates, fraction of multiply-mapping reads, number of unique 20-mers at the 5’ end of the reads, insert size distributions; fraction of ribosomal RNAs were calculated using bedtools v. 2.25.0. (Quinlan and Hall, 2010). In addition, each resulting bam file was randomly down-sampled to a million reads, which were aligned against hg19 and read density across genomic features were estimated for RNA-Seq-specific quality control metrics. RNA-Seq mapping and quantitation: Reads were aligned against GRCh38 / ENSEMBL 89 annotation using STAR v. 2.5.3a (Dobin et al., 2013) with the following flags -runThreadN 8 --runMode alignReads --outFilterType BySJout --outFilterMultimapNmax 20 --alignSJoverhangMin 8 --alignSJDBoverhangMin 1 -- outFilterMismatchNmax 999 --alignIntronMin 10 --alignIntronMax 1000000 --alignMatesGapMax 1000000 --outSAMtype BAM SortedByCoordinate --quantMode TranscriptomeSAM with -- genomeDir pointing to a 75nt-junction GRCh38 STAR suffix array.

Gene expression was quantitated using RSEM v. 1.3.0 (Li and Dewey, 2011) with the following flags for all libraries: rsem-calculate-expression --calc-pme --alignments -p 8 --forward-prob 0 against an annotation matching the STAR SA reference. Posterior mean estimates (pme) of counts and estimated RPKM were retrieved.

#### Differential gene expression analysis

To identify significantly altered genes in isolation vs interaction conditions, differential gene analysis of count data was performed using DESeq2 (Version 1.12.3) in R (Love et al., 2014). Dataset parameters were estimated using the estimateSizeFactors(), and estimateDispersions() functions; read counts across conditions were modeled based on a negative binomial distribution and a Wald test was used to test for differential expression (nbinomWaldtest(), all packaged into the DESeq() function), using the treatment type as a contrast. Fold-changes, *p*-values and Benjamin-Hochberg-adjusted *p*-values (BH) were reported for each protein-coding gene. Regularized fold-changes were calculated using the lfcShrink() function. Heat-maps were generated in Spotfire (Tibco), using a complete linkage function and the cosine correlation as a distance metrics to compute gene-level dendrograms and sample-level for the Cell marker heatmap. Clades from the gene-level hierarchical clustering were used to subset genes for gene expression time-course representations.

#### Gene set enrichment analysis, pathway enrichment, gene ontology analysis and visualization

Differential expression results from DESeq2 were retrieved, and the “stat” column was used to pre-rank genes for GSEA analysis. Briefly, the “stat” values reflect the Wald’s test performed on read counts as modeled by DESeq2 using the negative binomial distribution. Genes that weren’t expressed were excluded from the analysis. GSEA (version 2.2.3) was performed to identify differentially regulated gene sets in isolation versus interaction, as described previously (Subramanian et al., 2005). To stabilize variance, the normalized count data were processed using a regularized logarithm transformation in DESeq2 (Love et al., 2014). The signal-to-noise metric was used to generate the ranked list of genes. The empirical p-values for each enrichment score were calculated relative to the null distribution of enrichment scores, which was computed via 1000 gene set permutations. Gene sets with nominal p-value <0.05 and q-value <0.05 were considered significant. Volcano plots of differentially expressed genes were made with plot.ly (Plotly). Positive or negative fold changes of differential gene expressions were analyzed separately for enrichments. Pathway analysis as well as gene ontology term analysis based on various databases were performed using the following tools: Enrichr (Kuleshov et al., 2016) and ClueGO (Bindea et al., 2009) in Cytoscape v3.5.1 (Cytoscape consortium). ClueGo was also used for visualization of significantly enriched KEGG networks where size and color intensity of nods correspond to significance of enrichments. Venn diagrams were created with Venny 2.0 (Oliveros, 2007-2015).

#### RNA-Metabolomic integration

Correlation in positive enrichments of metabolism related genes of each MPS and the positive enrichments of metabolites in the common media was determined with Joint pathway analysis using MetaboAnalyst 4 (McGill university) (Chong et al., 2018). The results represent common changes in metabolic activity of individual MPS.

### Data Availability

The GEO accession number for the RNA-seq data reported in this paper is GSE135428.

Raw RNAseq data of the UC epithelial donor could not be included due to sharing restrictions however list of differentially expressed genes as quality contorols for the comparison among conditions can be shared upon request to the lead author.

## Notes

#### Summary of Updates

- List of Authors - Figures

## References

Adams, D.H., and Eksteen, B. (2006). Aberrant homing of mucosal T cells and extra-intestinal manifestations of inflammatory bowel disease. Nat Rev Immunol 6, 244–251.

Ananthakrishnan, A.N., Khalili, H., Konijeti, G.G., Higuchi, L.M., de Silva, P., Korzenik, J.R., Fuchs, C.S., Willett, W.C., Richter, J.M., and Chan, A.T. (2013). A prospective study of long-term intake of dietary fiber and risk of Crohn’s disease and ulcerative colitis. Gastroenterology 145, 970–977.

Barr, M., Gately, K., and O’Byrne, K. (2012). Vascular endothelial growth factor (VEGF) is an autocrine survival factor in non-small cell lung cancer. Lung Cancer 75, S2–S2.

Belyayev, L., Loh, K., Fishbein, T.M., and Kroemer, A. (2019). The parallel paradigm between intestinal transplant inflammation and inflammatory bowel disease. Curr Opin Organ Tran 24, 207–211.

Berndt, B.E., Zhang, M., Owyang, S.Y., Cole, T.S., Wang, T.W., Luther, J., Veniaminova, N.A., Merchant, J.L., Chen, C.C., Huffnagle, G.B., et al. (2012). Butyrate increases IL-23 production by stimulated dendritic cells. Am J Physiol Gastrointest Liver Physiol 303, G1384–1392.

Bindea, G., Mlecnik, B., Hackl, H., Charoentong, P., Tosolini, M., Kirilovsky, A., Fridman, W.H., Pages, F., Trajanoski, Z., and Galon, J. (2009). ClueGO: a Cytoscape plug-in to decipher functionally grouped gene ontology and pathway annotation networks. Bioinformatics 25, 1091–1093.

Breuer, R.I., Soergel, K.H., Lashner, B.A., Christ, M.L., Hanauer, S.B., Vanagunas, A., Harig, J.M., Keshavarzian, A., Robinson, M., Sellin, J.H., et al. (1997). Short chain fatty acid rectal irrigation for left-sided ulcerative colitis: a randomised, placebo controlled trial. Gut 40, 485–491.

Carbone, M., and Neuberger, J.M. (2014). Autoimmune liver disease, autoimmunity and liver transplantation. J Hepatol 60, 210–223.

Chang, P.V., Hao, L., Offermanns, S., and Medzhitov, R. (2014). The microbial metabolite butyrate regulates intestinal macrophage function via histone deacetylase inhibition. Proc Natl Acad Sci U S A 111, 2247–2252.

Chen, W.L.K., Edington, C., Suter, E., Yu, J., Velazquez, J.J., Velazquez, J.G., Shockley, M., Large, E.M., Venkataramanan, R., Hughes, D.J., et al. (2017). Integrated gut/liver microphysiological systems elucidates inflammatory inter-tissue crosstalk. Biotechnol Bioeng 114, 2648–2659.

Cholongitas, E., Shusang, V., Papatheodoridis, G.V., Marelli, L., Manousou, P., Rolando, N., Patch, D., Rolles, K., Davidson, B., and Burroughs, A.K. (2008). Risk factors for recurrence of primary sclerosing cholangitis after liver transplantation. Liver Transpl 14, 138–143.

Chong, J., Soufan, O., Li, C., Caraus, I., Li, S., Bourque, G., Wishart, D.S., and Xia, J. (2018). MetaboAnalyst 4.0: towards more transparent and integrative metabolomics analysis. Nucleic Acids Res 46, W486–W494.

Clark, A.M., Wheeler, S.E., Young, C.L., Stockdale, L., Shepard Neiman, J., Zhao, W., Stolz, D.B., Venkataramanan, R., Lauffenburger, D., Griffith, L., et al. (2016). A liver microphysiological system of tumor cell dormancy and inflammatory responsiveness is affected by scaffold properties. Lab Chip 17, 156–168.

Cummings, J.H., Pomare, E.W., Branch, W.J., Naylor, C.P., and Macfarlane, G.T. (1987). Short chain fatty acids in human large intestine, portal, hepatic and venous blood. Gut 28, 1221–1227.

DeRoche, T.C., Xiao, S.Y., and Liu, X. (2014). Histological evaluation in ulcerative colitis. Gastroenterol Rep (Oxf) 2, 178–192.

Desmet, V.J., Gerber, M., Hoofnagle, J.H., Manns, M., and Scheuer, P.J. (1994). Classification of chronic hepatitis: diagnosis, grading and staging. Hepatology 19, 1513–1520.

Dobin, A., Davis, C.A., Schlesinger, F., Drenkow, J., Zaleski, C., Jha, S., Batut, P., Chaisson, M., and Gingeras, T.R. (2013). STAR: ultrafast universal RNA-seq aligner. Bioinformatics 29, 15–21.

Eastaff-Leung, N., Mabarrack, N., Barbour, A., Cummins, A., and Barry, S. (2010). Foxp3+ regulatory T cells, Th17 effector cells, and cytokine environment in inflammatory bowel disease. J Clin Immunol 30, 80–89.

Edington, C.D., Chen, W.L.K., Geishecker, E., Kassis, T., Soenksen, L.R., Bhushan, B.M., Freake, D., Kirschner, J., Maass, C., Tsamandouras, N., et al. (2018). Interconnected Microphysiological Systems for Quantitative Biology and Pharmacology Studies. Sci Rep 8, 4530.

Eltzschig, H.K., Sitkovsky, M.V., and Robson, S.C. (2013). Purinergic signaling during inflammation. N Engl J Med 368, 1260.

Erben, U., Loddenkemper, C., Doerfel, K., Spieckermann, S., Haller, D., Heimesaat, M.M., Zeitz, M., Siegmund, B., and Kuhl, A.A. (2014). A guide to histomorphological evaluation of intestinal inflammation in mouse models. Int J Clin Exp Pathol 7, 4557–4576.

Evans AM, B.B., Liu Q, Mitchell MW, Robinson RJ, et al. (2014). High Resolution Mass Spectrometry Improves Data Quantity and Quality as Compared to Unit Mass Resolution Mass Spectrometry in High-Throughput Profiling Metabolomics. Metabolomics 4.

Feng, T.T., Zou, T., Wang, X., Zhao, W.F., and Qin, A.L. (2017). Clinical significance of changes in the Th17/Treg ratio in autoimmune liver disease. World J Gastroenterol 23, 3832–3838.

Furusawa, Y., Obata, Y., Fukuda, S., Endo, T.A., Nakato, G., Takahashi, D., Nakanishi, Y., Uetake, C., Kato, K., Kato, T., et al. (2013). Commensal microbe-derived butyrate induces the differentiation of colonic regulatory T cells. Nature 504, 446–450.

Fuss, I.J., Neurath, M., Boirivant, M., Klein, J.S., de la Motte, C., Strong, S.A., Fiocchi, C., and Strober, W. (1996). Disparate CD4+ lamina propria (LP) lymphokine secretion profiles in inflammatory bowel disease. Crohn’s disease LP cells manifest increased secretion of IFN-gamma, whereas ulcerative colitis LP cells manifest increased secretion of IL-5. J Immunol 157, 1261–1270.

Gieseck, R.L., 3rd, Wilson, M.S., and Wynn, T.A. (2018). Type 2 immunity in tissue repair and fibrosis. Nat Rev Immunol 18, 62–76.

Griffin, J.M., Gilbert, K.M., Lamps, L.W., and Pumford, N.R. (2000). CD4(+) T-cell activation and induction of autoimmune hepatitis following trichloroethylene treatment in MRL+/+ mice. Toxicol Sci 57, 345–352.

Henning, S.J., and Hird, F.J. (1972). Ketogenesis from butyrate and acetate by the caecum and the colon of rabbits. Biochem J 130, 785–790.

Hoving, J.C. (2018). Targeting IL-13 as a Host-Directed Therapy Against Ulcerative Colitis. Front Cell Infect Mi 8.

Ivanov, II, Atarashi, K., Manel, N., Brodie, E.L., Shima, T., Karaoz, U., Wei, D., Goldfarb, K.C., Santee, C.A., Lynch, S.V., et al. (2009). Induction of intestinal Th17 cells by segmented filamentous bacteria. Cell 139, 485–498.

Kaiko, G.E., Ryu, S.H., Koues, O.I., Collins, P.L., Solnica-Krezel, L., Pearce, E.J., Pearce, E.L., Oltz, E.M., and Stappenbeck, T.S. (2016). The Colonic Crypt Protects Stem Cells from Microbiota-Derived Metabolites. Cell 167, 1137.

Kasendra, M., Tovaglieri, A., Sontheimer-Phelps, A., Jalili-Firoozinezhad, S., Bein, A., Chalkiadaki, A., Scholl, W., Zhang, C., Rickner, H., Richmond, C.A., et al. (2018). Development of a primary human Small Intestine-on-a-Chip using biopsy-derived organoids. Sci Rep 8, 2871.

Kespohl, M., Vachharajani, N., Luu, M., Harb, H., Pautz, S., Wolff, S., Sillner, N., Walker, A., Schmitt-Kopplin, P., Boettger, T., et al. (2017). The Microbial Metabolite Butyrate Induces Expression of Th1-Associated Factors in CD4(+) T Cells. Front Immunol 8, 1036.

Khor, B., Gardet, A., and Xavier, R.J. (2011). Genetics and pathogenesis of inflammatory bowel disease. Nature 474, 307–317.

Kim, M.H., Kang, S.G., Park, J.H., Yanagisawa, M., and Kim, C.H. (2013). Short-chain fatty acids activate GPR41 and GPR43 on intestinal epithelial cells to promote inflammatory responses in mice. Gastroenterology 145, 396–406 e391-310.

Kimura, I., Inoue, D., Maeda, T., Hara, T., Ichimura, A., Miyauchi, S., Kobayashi, M., Hirasawa, A., and Tsujimoto, G. (2011). Short-chain fatty acids and ketones directly regulate sympathetic nervous system via G protein-coupled receptor 41 (GPR41). Proc Natl Acad Sci U S A 108, 8030–8035.

Kinchen, J., Chen, H.H., Parikh, K., Antanaviciute, A., Jagielowicz, M., Fawkner-Corbett, D., Ashley, N., Cubitt, L., Mellado-Gomez, E., Attar, M., et al. (2018). Structural Remodeling of the Human Colonic Mesenchyme in Inflammatory Bowel Disease. Cell 175, 372-+.

King, C.G., Kobayashi, T., Cejas, P.J., Kim, T., Yoon, K., Kim, G.K., Chiffoleau, E., Hickman, S.P., Walsh, P.T., Turka, L.A., et al. (2006). TRAF6 is a T cell-intrinsic negative regulator required for the maintenance of immune homeostasis. Nat Med 12, 1088–1092.

Kleiner, G., Zanin, V., Monasta, L., Crovella, S., Caruso, L., Milani, D., and Marcuzzi, A. (2015). Pediatric patients with inflammatory bowel disease exhibit increased serum levels of proinflammatory cytokines and chemokines, but decreased circulating levels of macrophage inhibitory protein-1beta, interleukin-2 and interleukin-17. Exp Ther Med 9, 2047–2052.

Koh, A., De Vadder, F., Kovatcheva-Datchary, P., and Backhed, F. (2016). From Dietary Fiber to Host Physiology: Short-Chain Fatty Acids as Key Bacterial Metabolites. Cell 165, 1332–1345.

Kuleshov, M.V., Jones, M.R., Rouillard, A.D., Fernandez, N.F., Duan, Q., Wang, Z., Koplev, S., Jenkins, S.L., Jagodnik, K.M., Lachmann, A., et al. (2016). Enrichr: a comprehensive gene set enrichment analysis web server 2016 update. Nucleic Acids Res 44, W90–97.

Lavoie, S., Conway, K.L., Lassen, K.G., Jijon, H.B., Pan, H., Chun, E., Michaud, M., Lang, J.K., Gallini Comeau, C.A., Dreyfuss, J.M., et al. (2019). The Crohn’s disease polymorphism, ATG16L1 T300A, alters the gut microbiota and enhances the local Th1/Th17 response. Elife 8.

Li, B., and Dewey, C.N. (2011). RSEM: accurate transcript quantification from RNA-Seq data with or without a reference genome. BMC Bioinformatics 12, 323.

Lian, J.S., Liu, W., Hao, S.R., Chen, D.Y., Wang, Y.Y., Yang, J.L., Jia, H.Y., and Huang, J.R. (2015). A serum metabolomic analysis for diagnosis and biomarker discovery of primary biliary cirrhosis and autoimmune hepatitis. Hepatobiliary Pancreat Dis Int 14, 413–421.

Liu, Y., Meyer, C., Xu, C., Weng, H., Hellerbrand, C., ten Dijke, P., and Dooley, S. (2013). Animal models of chronic liver diseases. Am J Physiol Gastrointest Liver Physiol 304, G449–468.

Loftus, E.V., Jr., Harewood, G.C., Loftus, C.G., Tremaine, W.J., Harmsen, W.S., Zinsmeister, A.R., Jewell, D.A., and Sandborn, W.J. (2005). PSC-IBD: a unique form of inflammatory bowel disease associated with primary sclerosing cholangitis. Gut 54, 91–96.

Long, T.J., Cosgrove, P.A., Dunn, R.T., 2nd, Stolz, D.B., Hamadeh, H., Afshari, C., McBride, H., and Griffith, L.G. (2016). Modeling Therapeutic Antibody-Small Molecule Drug-Drug Interactions Using a Three-Dimensional Perfusable Human Liver Coculture Platform. Drug Metab Dispos 44, 1940–1948.

Love, M.I., Huber, W., and Anders, S. (2014). Moderated estimation of fold change and dispersion for RNA-seq data with DESeq2. Genome Biol 15, 550.

Luu, M., Weigand, K., Wedi, F., Breidenbend, C., Leister, H., Pautz, S., Adhikary, T., and Visekruna, A. (2018). Regulation of the effector function of CD8(+) T cells by gut microbiota-derived metabolite butyrate. Sci Rep-Uk 8.

Macdonald, T.T., and Monteleone, G. (2005). Immunity, inflammation, and allergy in the gut. Science 307, 1920–1925.

Maeda, M., Watanabe, N., Neda, H., Yamauchi, N., Okamoto, T., Sasaki, H., Tsuji, Y., Akiyama, S., Tsuji, N., and Niitsu, Y. (1992). Serum tumor necrosis factor activity in inflammatory bowel disease. Immunopharmacol Immunotoxicol 14, 451–461.

Metsalu, T., and Vilo, J. (2015). ClustVis: a web tool for visualizing clustering of multivariate data using Principal Component Analysis and heatmap. Nucleic Acids Res 43, W566–570.

Morrison, D.J., and Preston, T. (2016). Formation of short chain fatty acids by the gut microbiota and their impact on human metabolism. Gut Microbes 7, 189–200.

Murai, M., Yoneyama, H., Ezaki, T., Suematsu, M., Terashima, Y., Harada, A., Hamada, H., Asakura, H., Ishikawa, H., and Matsushima, K. (2003). Peyer’s patch is the essential site in initiating murine acute and lethal graft-versus-host reaction. Nat Immunol 4, 154–160.

O’Keefe, S.J.D. (2016). Diet, microorganisms and their metabolites, and colon cancer. Nat Rev Gastro Hepat 13, 691–706.

Okin, D., and Medzhitov, R. (2016). The Effect of Sustained Inflammation on Hepatic Mevalonate Pathway Results in Hyperglycemia. Cell 165, 343–356.

Oliveros, J.C. (2007-2015). Venny. An interactive tool for comparing lists with Venn’s diagrams.

Park, J., Goergen, C.J., HogenEsch, H., and Kim, C.H. (2016). Chronically Elevated Levels of Short-Chain Fatty Acids Induce T Cell-Mediated Ureteritis and Hydronephrosis. J Immunol 196, 2388–2400.

Park, J., Kim, M., Kang, S.G., Jannasch, A.H., Cooper, B., Patterson, J., and Kim, C.H. (2015). Short-chain fatty acids induce both effector and regulatory T cells by suppression of histone deacetylases and regulation of the mTOR-S6K pathway. Mucosal Immunology 8, 80–93.

Perdigoto, R., Carpenter, H.A., and Czaja, A.J. (1992). Frequency and significance of chronic ulcerative colitis in severe corticosteroid-treated autoimmune hepatitis. J Hepatol 14, 325–331.

Podolsky, D.K. (1991). Inflammatory bowel disease (2). N Engl J Med 325, 1008–1016.

Puchalska, P., and Crawford, P.A. (2017). Multi-dimensional Roles of Ketone Bodies in Fuel Metabolism, Signaling, and Therapeutics. Cell Metab 25, 262–284.

Quinlan, A.R., and Hall, I.M. (2010). BEDTools: a flexible suite of utilities for comparing genomic features. Bioinformatics 26, 841–842.

Ratajczak, P., Janin, A., Peffault de Latour, R., Leboeuf, C., Desveaux, A., Keyvanfar, K., Robin, M., Clave, E., Douay, C., Quinquenel, A., et al. (2010). Th17/Treg ratio in human graft-versus-host disease. Blood 116, 1165–1171.

Ronaldson-Bouchard, K., and Vunjak-Novakovic, G. (2018). Organs-on-a-Chip: A Fast Track for Engineered Human Tissues in Drug Development. Cell Stem Cell 22, 310–324.

Roper, J., Tammela, T., Cetinbas, N.M., Akkad, A., Roghanian, A., Rickelt, S., Almeqdadi, M., Wu, K., Oberli, M.A., Sanchez-Rivera, F.J., et al. (2017). In vivo genome editing and organoid transplantation models of colorectal cancer and metastasis. Nat Biotechnol 35, 569–576.

Roth, E., Oehler, R., Manhart, N., Exner, R., Wessner, B., Strasser, E., and Spittler, A. (2002). Regulative potential of glutamine - Relation to glutathione metabolism. Nutrition 18, 217–221.

Rudensky, A.Y. (2011). Regulatory T cells and Foxp3. Immunol Rev 241, 260–268.

Sampson, T.R., Debelius, J.W., Thron, T., Janssen, S., Shastri, G.G., Ilhan, Z.E., Challis, C., Schretter, C.E., Rocha, S., Gradinaru, V., et al. (2016). Gut Microbiota Regulate Motor Deficits and Neuroinflammation in a Model of Parkinson’s Disease. Cell 167, 1469–1480 e1412.

Sarkar, U., Rivera-Burgos, D., Large, E.M., Hughes, D.J., Ravindra, K.C., Dyer, R.L., Ebrahimkhani, M.R., Wishnok, J.S., Griffith, L.G., and Tannenbaum, S.R. (2015). Metabolite profiling and pharmacokinetic evaluation of hydrocortisone in a perfused three-dimensional human liver bioreactor. Drug Metab Dispos 43, 1091–1099.

Smith, P.M., Howitt, M.R., Panikov, N., Michaud, M., Gallini, C.A., Bohlooly, Y.M., Glickman, J.N., and Garrett, W.S. (2013). The microbial metabolites, short-chain fatty acids, regulate colonic Treg cell homeostasis. Science 341, 569–573.

Srinivasan, B., Kolli, A.R., Esch, M.B., Abaci, H.E., Shuler, M.L., and Hickman, J.J. (2015). TEER measurement techniques for in vitro barrier model systems. J Lab Autom 20, 107–126.

Subramanian, A., Tamayo, P., Mootha, V.K., Mukherjee, S., Ebert, B.L., Gillette, M.A., Paulovich, A., Pomeroy, S.L., Golub, T.R., Lander, E.S., et al. (2005). Gene set enrichment analysis: a knowledge-based approach for interpreting genome-wide expression profiles. Proc Natl Acad Sci U S A 102, 15545–15550.

Taneja, V., and David, C.S. (2001). Lessons from animal models for human autoimmune diseases. Nat Immunol 2, 781–784.

Tarrerias, A.L., Millecamps, M., Alloui, A., Beaughard, C., Kemeny, J.L., Bourdu, S., Bommelaer, G., Eschalier, A., Dapoigny, M., and Ardid, D. (2002). Short-chain fatty acid enemas fail to decrease colonic hypersensitivity and inflammation in TNBS-induced colonic inflammation in rats. Pain 100, 91–97.

Trompette, A., Gollwitzer, E.S., Pattaroni, C., Lopez-Mejia, I.C., Riva, E., Pernot, J., Ubags, N., Fajas, L., Nicod, L.P., and Marsland, B.J. (2018). Dietary Fiber Confers Protection against Flu by Shaping Ly6c(-) Patrolling Monocyte Hematopoiesis and CD8(+) T Cell Metabolism. Immunity 48, 992–1005 e1008.

Tsamandouras, N., Chen, W.L.K., Edington, C.D., Stokes, C.L., Griffith, L.G., and Cirit, M. (2017). Integrated Gut and Liver Microphysiological Systems for Quantitative In Vitro Pharmacokinetic Studies. AAPS J 19, 1499–1512.

Wang, A., Luan, H.H., and Medzhitov, R. (2019). An evolutionary perspective on immunometabolism. Science 363.

West, N.P., Christophersen, C.T., Pyne, D.B., Cripps, A.W., Conlon, M.A., Topping, D.L., Kang, S., McSweeney, C.S., Fricker, P.A., Aguirre, D., et al. (2013). Butyrylated starch increases colonic butyrate concentration but has limited effects on immunity in healthy physically active individuals. Exerc Immunol Rev 19, 102–119.

Wu, J., Zhang, H., Shi, X., Xiao, X., Fan, Y., Minze, L.J., Wang, J., Ghobrial, R.M., Xia, J., Sciammas, R., et al. (2017). Ablation of Transcription Factor IRF4 Promotes Transplant Acceptance by Driving Allogenic CD4(+) T Cell Dysfunction. Immunity 47, 1114–1128 e1116.

Yan, G., Li, L., Zhu, B., and Li, Y. (2016). Lipidome in colorectal cancer. Oncotarget 7, 33429–33439.

Yissachar, N., Zhou, Y., Ung, L., Lai, N.Y., Mohan, J.F., Ehrlicher, A., Weitz, D.A., Kasper, D.L., Chiu, I.M., Mathis, D., et al. (2017). An Intestinal Organ Culture System Uncovers a Role for the Nervous System in Microbe-Immune Crosstalk. Cell 168, 1135–1148 e1112.

Zhou, L., and Sonnenberg, G.F. (2018). Essential immunologic orchestrators of intestinal homeostasis. Sci Immunol 3.

