## Supplementary material for "Gut-Liver physiomimetics reveal paradoxical modulation of IBD-related inflammation by short-chain fatty acids"

**This PDF file includes:**

Figs. S1 to S7

### Supplementary figures

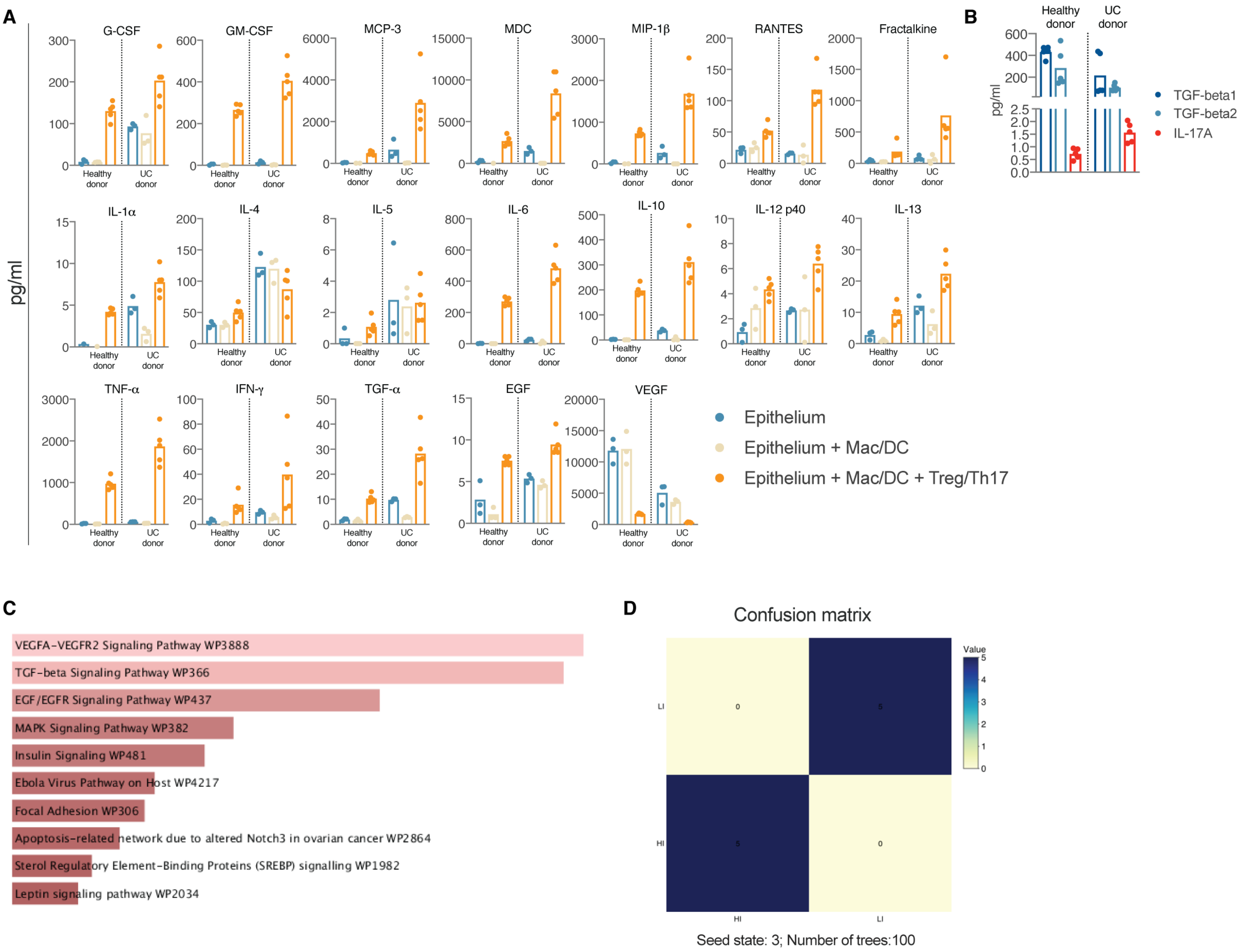


**Fig.S1**

**Gut donor of each MPS can be predicted with 100% accuracy by random forest analysis based on cytokines released during gut MPS-Treg/Th17 interaction.** (A) Representative values of individual cytokines measured in media of UC and healthy epithelial monolayers, UC and healthy gut MPS and MPS co-cultures with Treg/Th17 cells (B) Concentration of TGF-β1, TGF-β2 and IL-17 in basal media of UC and helthy gut MPS during interaction with Treg/Th17 cells (C) Wikipathway analysis of enriched pathways in UC gut MPS over healthy control (D) Random forest analysis of cytokines shown under (A) during UC and helthy MPS interaction with Treg/Th17 cells. Confusion matrix indicates prediction accuracy. Each condition was tested in separate experiments with 3-5 biological replicates.

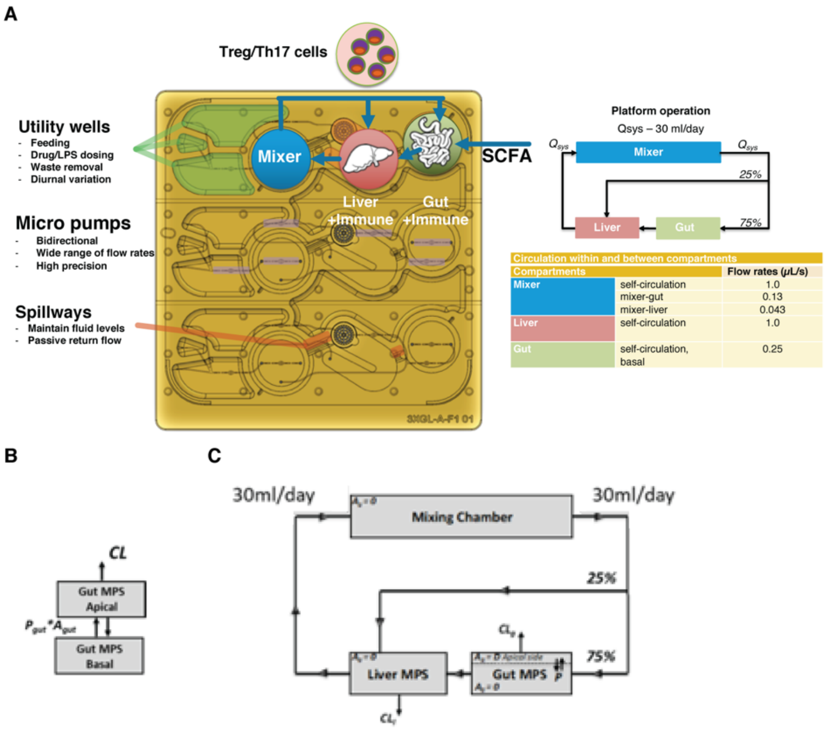


**Fig.S2**

**Platform operation, circulating CD4 T cell viability and schematic models to calculate SCFA metabolism and distribution.** (A) Operational parameters of the 3XGL platform with the gut and liver MPSs. (B) Schematic overview of the utilized compartmental model to describe SCFA distribution, metabolism in the gut MPS; $P_{gut}$ = permeability, $A_{gut}$ = surface area of transwell, CL = metabolism. (C) Schematic overview of the four-comparmental model consisting of a gut, liver and mixing chamber MPS. In addition to gut-specific SCFA permeability and consumption, liver metabolism was implemented to describe the distribution.

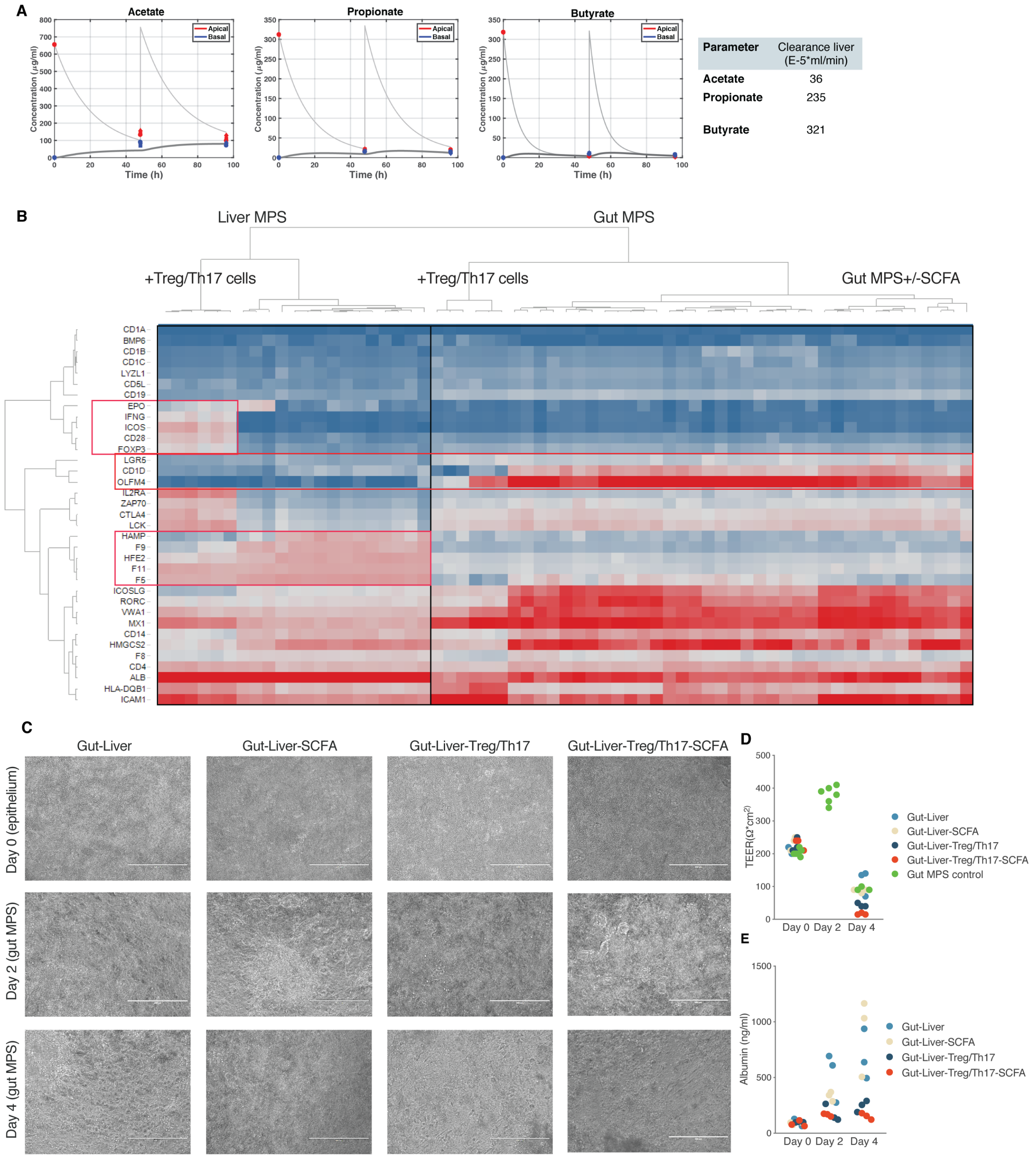


| Parameter | Clearance liver (E-5*ml/min) |
|---|---|
| Acetate | 36 |
| Propionate | 235 |
| Butyrate | 321 |

**Fig.S3**

**SCFA distribution and its effect on global gene expression and functional parameters of gut and liver function during gut-liver interaction with or without SCFA and Treg/Th17 cells.** (A) Apical concentrations of acetate, propionate and butyrate in UC gut MPS and their bioavailable concentrations in basal common medium during 96h of interaction. Far right: hepatic clearance of bioavailable SCFA. (B) Complete linkage and cluster heatmap of genes expressed by all tissues collected during the study.

(C) Representative brightfield images of epithelial monolayers at day 0 of interaction, prior to basolateral seeding of MACs/DCs, and that of gut MPS monolayers with underlying MACs/DCs at day 2 and day 4 of interaction (D) TEER of UC gut MPS used in the interaction studies as well as the control. D0: at the beginning of the interaction, D2: TEER of the control gut MPS in isolation, D4: end of the interaction. (E) Albumin concentrations produced by the liver MPS before, during and at the end of interaction measured in common medium.

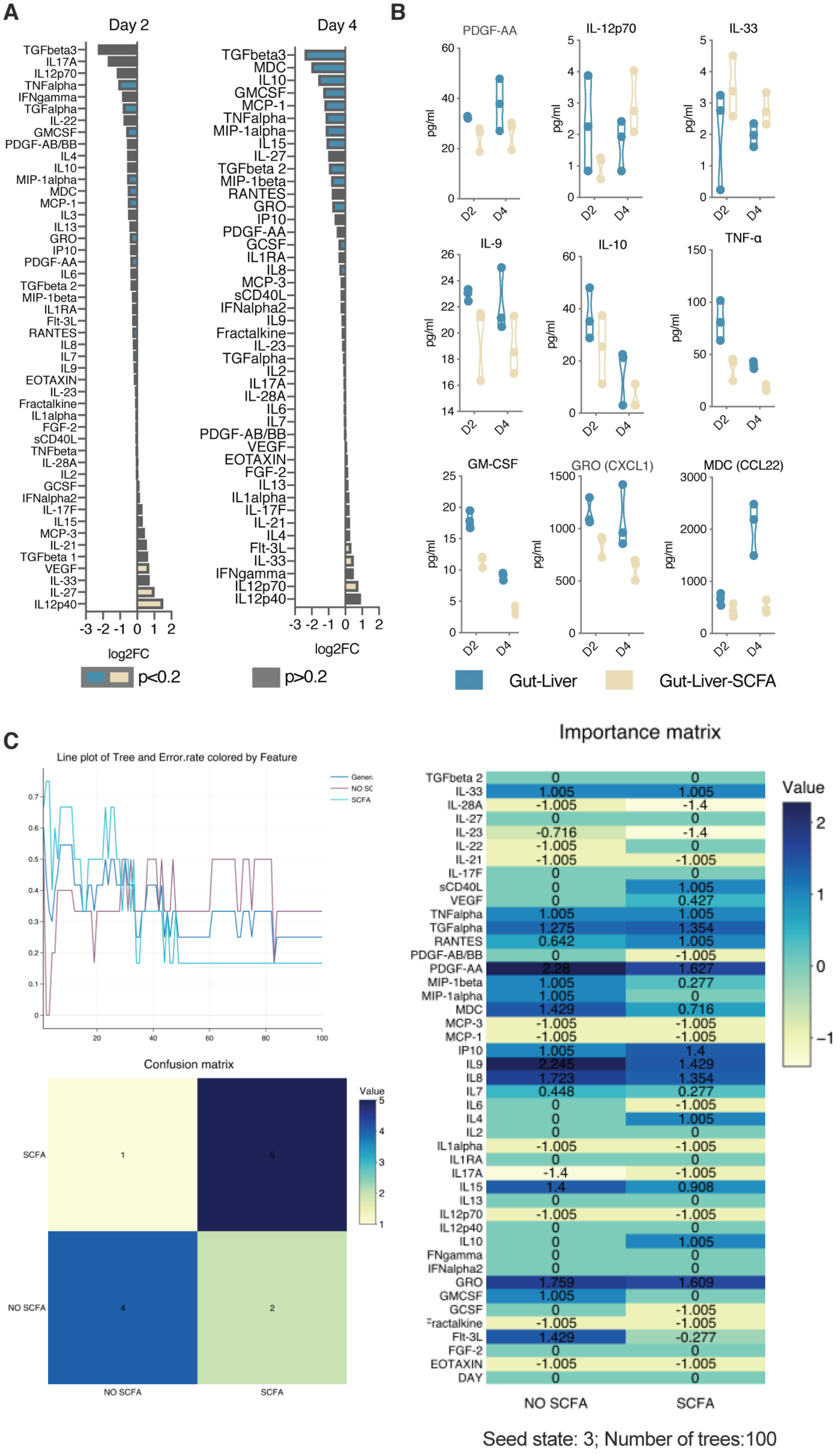
A
Day 2
Day 4
log2FC
p<0.2
p>0.2
B
PDGF-AA
IL-12p70
IL-33
IL-9
IL-10
TNF-α
GM-CSF
GRO (CXCL1)
MDC (CCL22)
pg/ml
D2
D4
Gut-Liver
Gut-Liver-SCFA
C
Line plot of Tree and Error.rate colored by Feature
Confusion matrix
SCFA
NO SCFA
Value
Importance matrix
TGFbeta 2 0 0
IL-33 1.005 1.005
IL-28A -1.005 -1.4
IL-27 0 0
IL-23 -0.716 -1.4
IL-22 -1.005 0
IL-21 -1.005 -1.005
IL-17F 0 0
sCD40L 0 1.005
VEGF 0 0.427
TNFalpha 1.005 1.005
TGFalpha 1.275 1.354
RANTES 0.642 1.005
PDGF-AB/BB 0 -1.005
PDGF-AA 2.28 1.627
MIP-1beta 1.005 0.277
MIP-1alpha 1.005 0
MDC 1.429 0.716
MCP-3 -1.005 -1.005
MCP-1 -1.005 -1.005
IP10 1.005 1.4
IL9 2.245 1.429
IL8 1.723 1.354
IL7 0.448 0.277
IL6 0 -1.005
IL4 0 1.005
IL2 0 0
IL1alpha -1.005 -1.005
IL1RA 0 0
IL17A -1.4 -1.005
IL15 1.4 0.908
IL13 0 0
IL12p70 -1.005 -1.005
IL12p40 0 0
IL10 0 1.005
FNgamma 0 0
IFNalpha2 0 0
GRO 1.759 1.609
GMCSF 1.005 0
GCSF 0 -1.005
Fractalkine -1.005 -1.005
Flt-3L 1.429 -0.277
FGF-2 0 0
EOTAXIN -1.005 -1.005
DAY 0 0
NO SCFA
SCFA
Value
Seed state: 3; Number of trees:100

**Fig.S4**

**Effect of SCFA on cytokine/chemokine expression during gut-liver interaction in the absence of Treg//Th17 cells.** (A) log2 Fold changes of individual multiplexed cytokines/chemokines measured in the common medium of the gut-liver and gut-liver-SCFA interactions. Grey colored bars have a p-value over 0.2 and blue (gut-liver over gut-liver-SCFA) or beige colored (gut-liver-SCFA over gut-liver) bars have a p-value less than 0.2. (B) Concentrations of cytokines/chemokines identified by random forest analysis to be most predictive of condition. (C) Random forest analysis of all cytokines/chemokines measured. Top left: Number of trees required to complete confusion matrix. Bottom left: Confusion matrix indicating prediction accuracy. Right: Importance matrix of most predictive parameters.

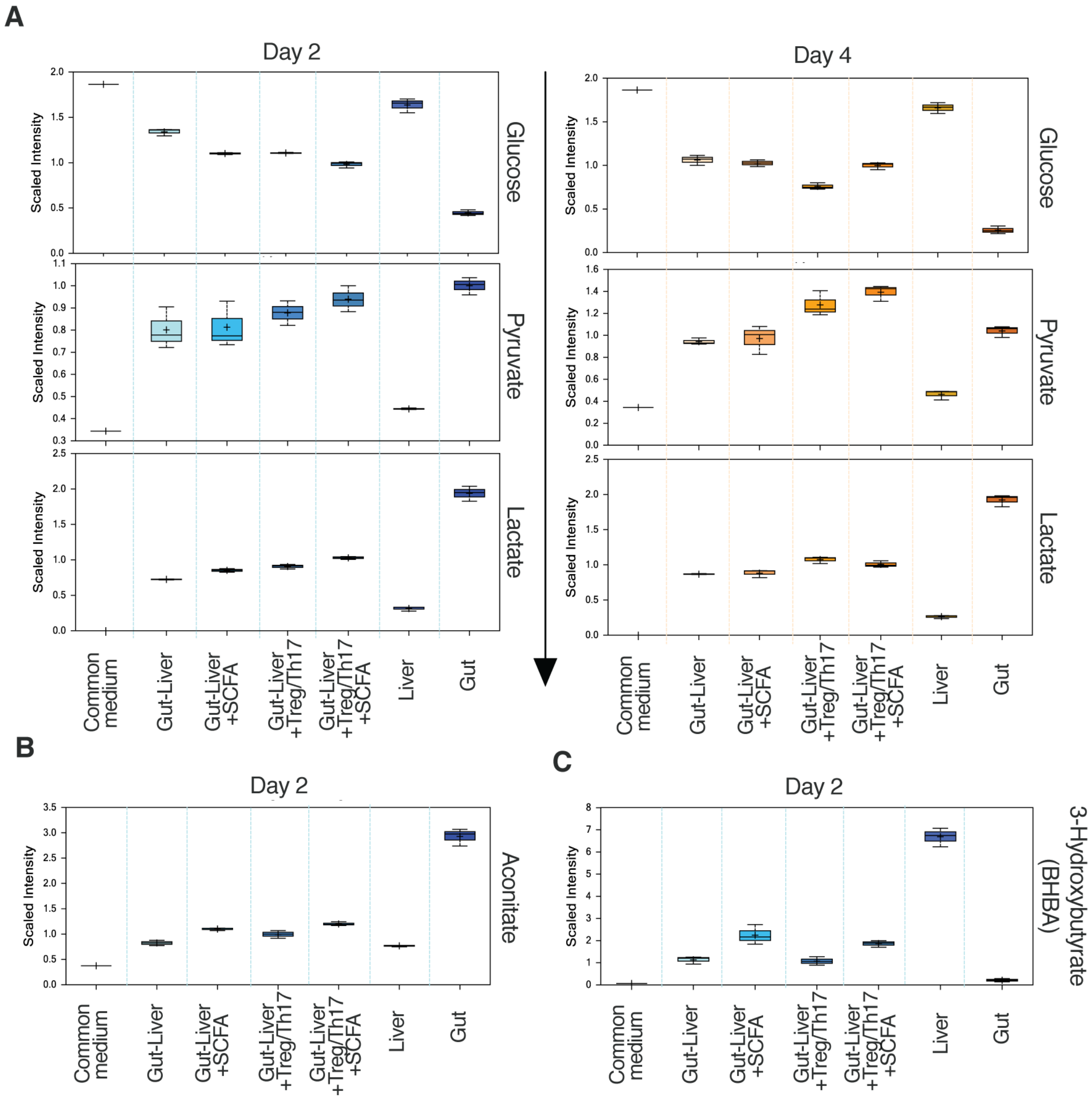


**Fig.S5**
**SCFA and systemic inflammation affect glucose metabolism and production of ketone bodies.** (A-C) Scaled intensities of glucose, pyruvate and lactate at days 2 and 4 (A), aconitate (B), and BHBA (C) at day 2 for interactions among all 4 experimental conditions, MPS controls in isolation and common media.

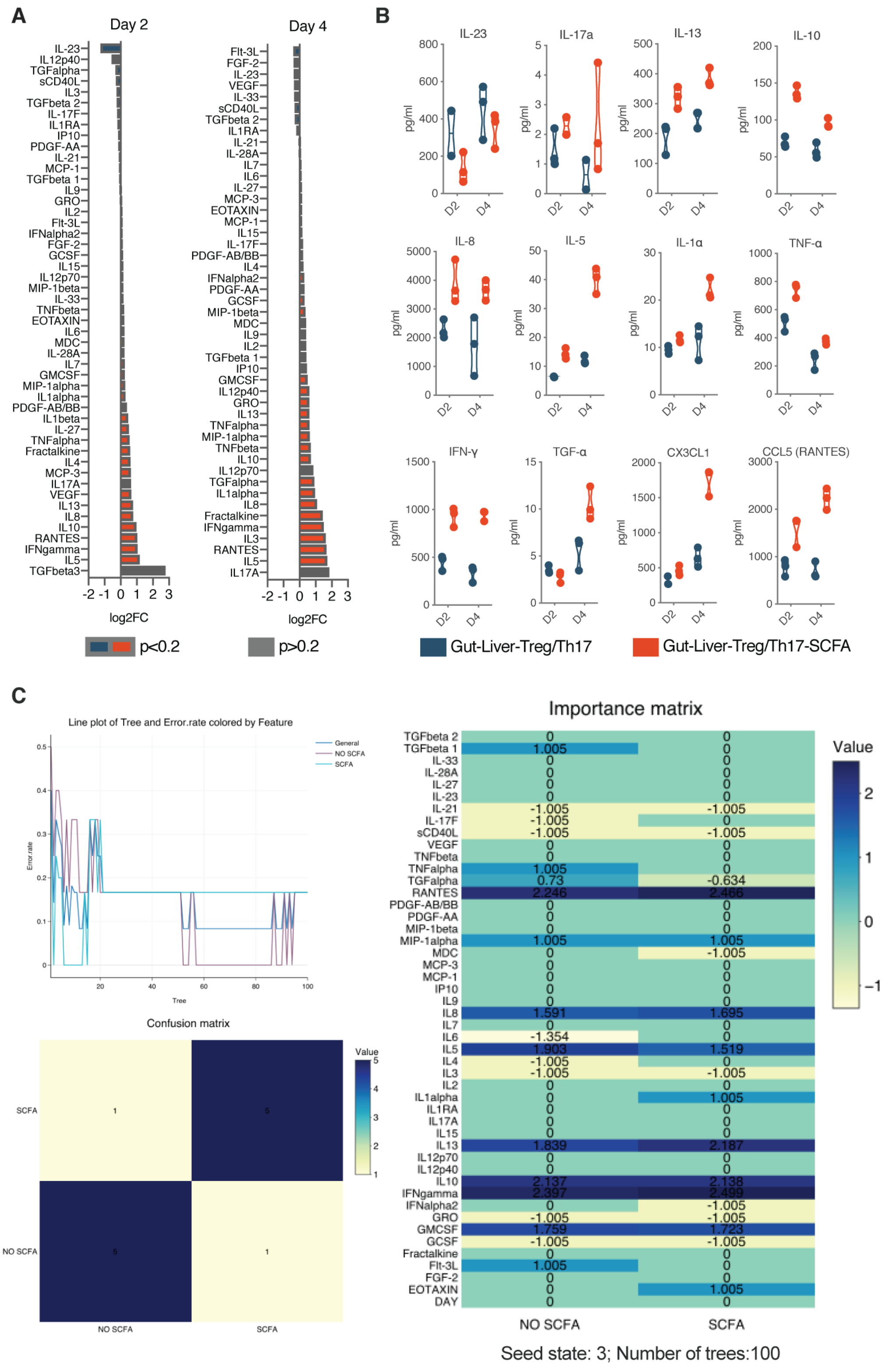
A
Day 2
Day 4
log2FC
p<0.2
p>0.2
B
IL-23
IL-17a
IL-13
IL-10
IL-8
IL-5
IL-1α
TNF-α
IFN-γ
TGF-α
CX3CL1
CCL5 (RANTES)
pg/ml
D2
D4
Gut-Liver-Treg/Th17
Gut-Liver-Treg/Th17-SCFA
C
Line plot of Tree and Error.rate colored by Feature
General
NO SCFA
SCFA
Error.rate
Tree
Confusion matrix
Value
NO SCFA
SCFA
Importance matrix
Value
NO SCFA
SCFA
Seed state: 3; Number of trees:100

**Fig.S6**

**Effect of SCFA on cytokine/chemokine expression during gut-liver interaction in the presence of Treg//Th17 cells** (A) log2 Fold changes of individual multiplexed cytokines/chemokines measured in the common media of the gut-liver-Treg/Th17 and gut-liver-Treg/Th17-SCFA interactions. Grey colored bars have a p-value over 0.2, and blue (gut-liver-Treg/Th17 over gut-liver-Treg/Th17-SCFA) or orange colored (gut-liver-Treg/Th17-SCFA over gut-liver-Treg/Th17) bars have a p-value less than 0.2. (B) Concentrations of cytokines/chemokines identified by random forest analysis to be most predictive of condition. (C) Random forest analysis of all cytokines/chemokines measured. Top left: Number of trees required to complete confusion matrix. Bottom left: Confusion matrix indicating prediction accuracy. Right: Importance matrix of most predictive parameters.

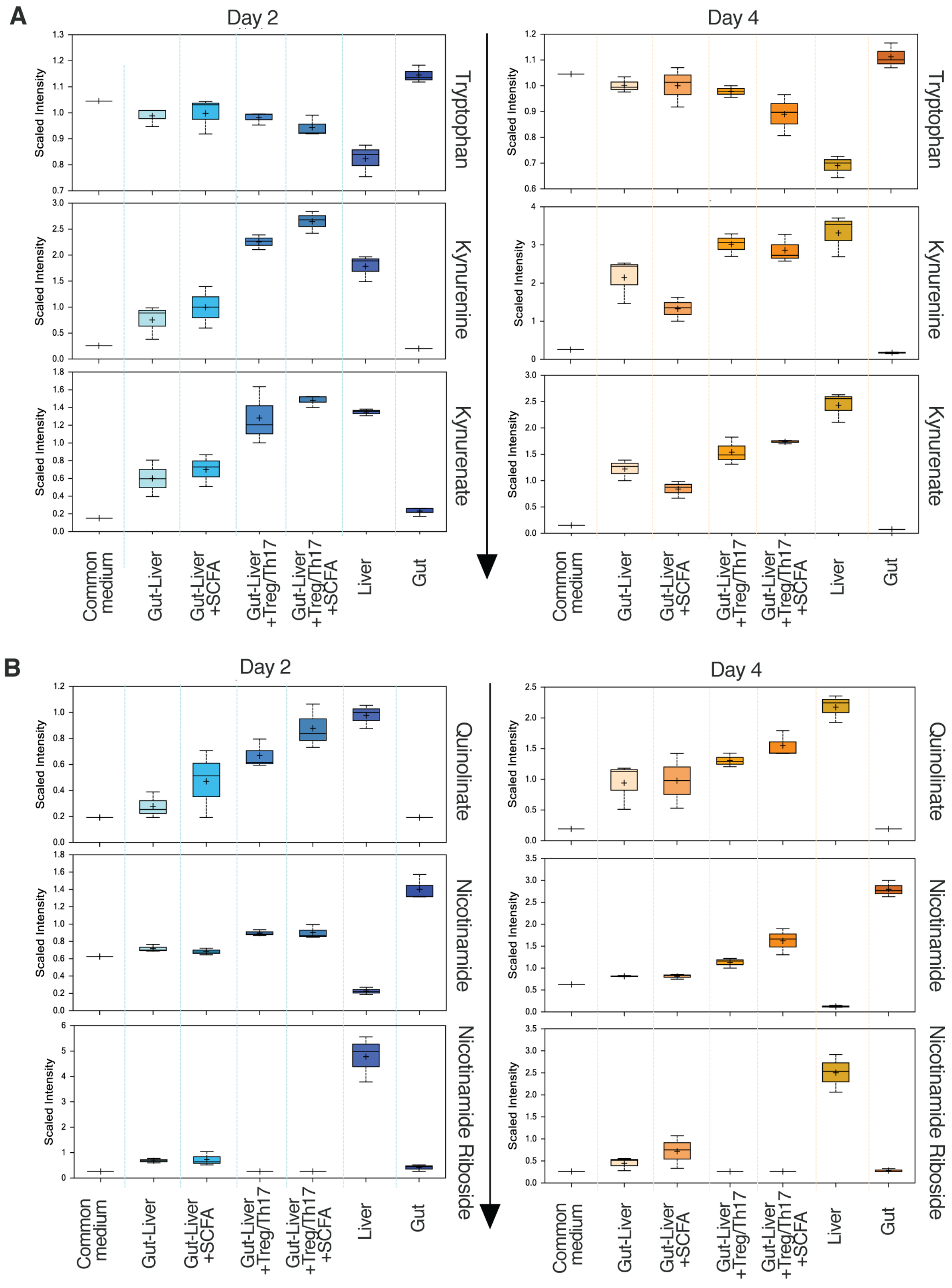


**Fig.S7**
**Effect of SCFA on inflammation related metabolic products during gut-liver interaction** (A,B) Scaled intensities of tryptophan, kynurenine and kynurenate (A) as well as quinolinate, nicotinamide and nicotinamide riboside (B) at days 2 and 4 for interactions among all 4 conditions, MPS controls in isolation and common media.
